# Multiplexed electron microscopy by fluorescent barcoding allows screening for ultrastructural phenotype

**DOI:** 10.1101/515841

**Authors:** Yury S. Bykov, Nir Cohen, Natalia Gabrielli, Hetty Manenschijn, Sonja Welsch, Petr Chlanda, Wanda Kukulski, Kiran R. Patil, Maya Schuldiner, John A.G. Briggs

## Abstract

Genetic screens performed using high-throughput fluorescent microscopes have generated large datasets that have contributed many insights into cell biology. However, such approaches typically cannot tackle questions requiring knowledge of ultrastructure below the resolution limit of fluorescent microscopy. Electron microscopy (EM) is not subject to this resolution limit, generating detailed images of cellular ultrastructure, but requires time consuming preparation of individual samples, limiting its throughput. Here we overcome this obstacle and describe a robust method for screening by high-throughput electron microscopy. Our approach uses combinations of fluorophores as barcodes to mark the genotype of each cell in mixed populations, and correlative light and electron microscopy to read the fluorescent barcode of each cell before it is imaged by electron microscopy. Coupled with an easy-to-use software workflow for correlation, segmentation and computer image analysis, our method allows to extract and analyze multiple cell populations from each EM sample preparation. We demonstrate the method on several organelles with samples that each contain up to 15 different yeast variants. The methodology is not restricted to yeast, can be scaled to higher-throughput, and can be utilized in multiple ways to enable electron microscopy to become a powerful screening methodology.

## Introduction

Functional studies can be extended from individual proteins to the whole-genomic level using high content screens relying on genetic tools, fluorescent light microscopy (LM), and automated workflows. Budding yeast (*Saccharomyces cerevisiae*, hereafter referred to simply as yeast) is a widely used model organism for high-throughput studies. Easy and scalable genetic manipulation has allowed the creation of tools such as systematic deletion libraries and GFP collections (Giaever et al., 2002; Yofe et al., 2016; Weill et al., 2018; Huh et al., 2003). Combined with automatic fluorescence microscopy these systematic libraries help to address a large variety of questions in cell biology (Ohya et al., 2005; Cohen and Schuldiner, 2011; Breker et al., 2014). Some questions cannot be addressed nor solved at the resolution limit of LM, but require higher resolution techniques such as electron microscopy (EM). However, EM has suffered, until now, from very low throughput.

With the introduction of fully computer-controlled electron microscopes, it is now possible to perform automated and large-scale data collection. This has particularly benefited the field of 3D EM which relies on the collection of a large number of projections (tomography) or serial sections (Mastronarde, 2005; Peddie and Collinson, 2014; Suloway et al., 2005; Zheng et al., 2004). The number of individual samples that can be analyzed by EM, however, remains relatively low because sample preparation procedures include time-consuming manual steps, and because samples are typically inserted individually into the electron microscope. To obtain the best preservation of both ultrastructure and fluorescence in yeast, each sample is subjected to high-pressure cryo-fixation, freeze-substitution, manual sectioning using a microtome, mounting on EM grids, insertion into the electron microscope and, of course, visualization and image analysis (McDonald, 2007; McDonald et al., 2010). These manual procedures have prevented applying the high-throughput screening paradigms that are common in LM, to the ultrastructural features that can be observed only by EM.

Here, we present an in-resin correlative light and electron microscopy (CLEM) approach (Spiegelhalter et al., 2014; Bykov et al., 2016; Kukulski et al., 2011) to increase the throughput of EM experiments (Fig. 1A). We apply it to yeast, but the approach can be adapted to other cell types. First, cells from different strains, genetic backgrounds or under different experimental conditions are grown in parallel. Then, each of them is treated by a unique combination of fluorescent labels creating a barcode. Following labeling, cells from parallel experiments are mixed together and undergo a single, unified, EM sample preparation. Prior to EM, the sample is imaged by LM to obtain the barcodes (staining patterns) for each cell. Correlation is performed between fluorescent and EM images, and cell positions and identities are determined. By multiplexing both sample preparation and EM imaging this method substantially increases potential throughput. Further, it removes the variability inherent in EM preparation of separate samples, allowing direct, automated comparison of images from parallel experiments and accurate quantification of traits.

**Fig. 1.**
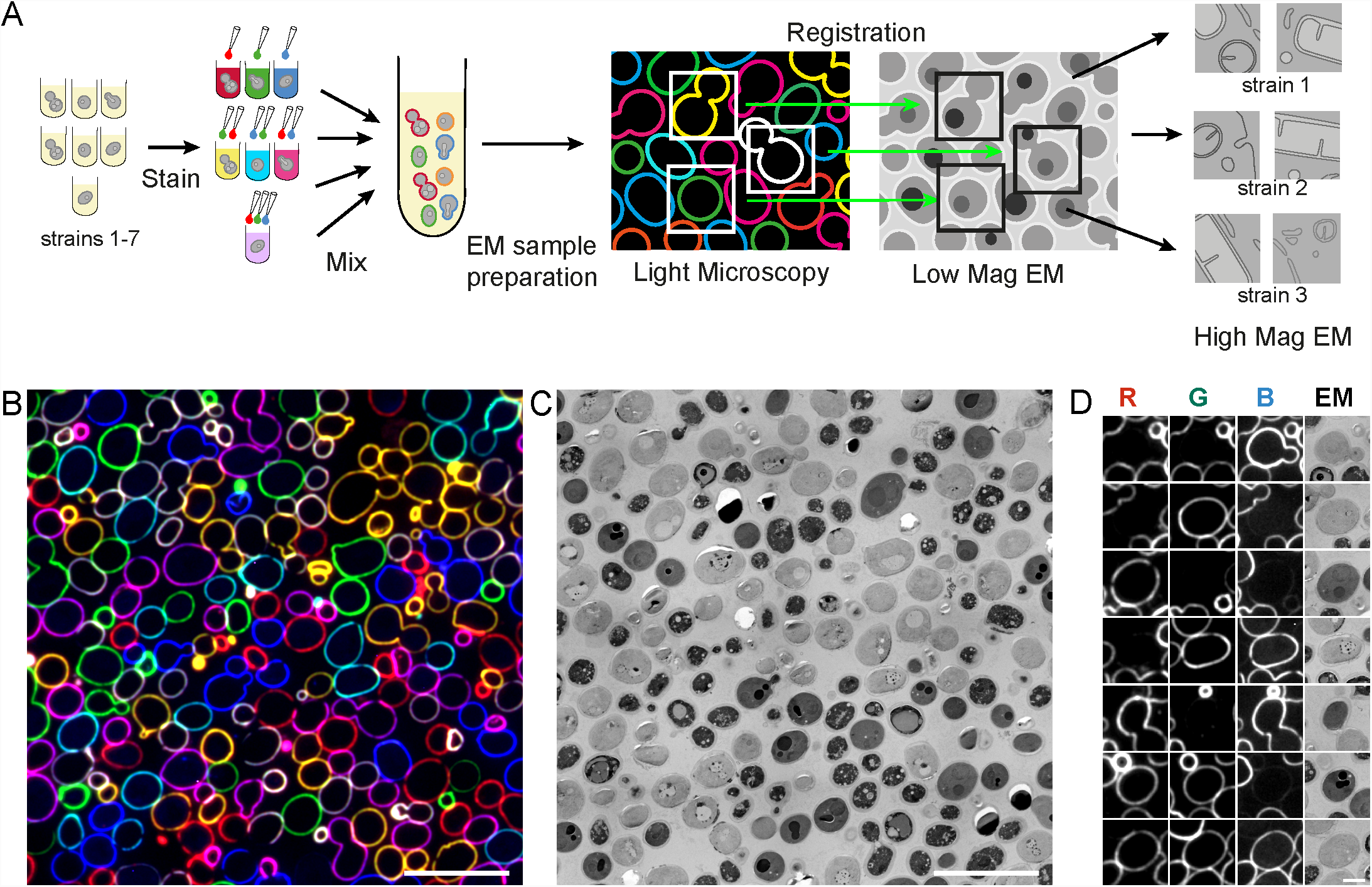
The principle of fluorescent barcoding CLEM. (A) Workflow schematic: strains or cells in different experimental conditions are grown in parallel and each tube is stained with a unique combina-tions of Con A conjugates (figure demonstrates three colors but more can be used) to produce a combi-nation (the barcode); then strains are mixed together and processed for EM. Samples are imaged in the LM and EM at medium magnification, correlation is performed, cell identities are determined using the fluorescent barcode, and the high-resolution information is collected for each strain. (B) Example using three-fluorophores to give seven well-discriminated combinations, pseudocolor composite image of a 100 nm thick Lowicryl section of embedded cells labeled with Con A conjugated to Alexa Fluor 350, Alexa Fluor 488, and TRITC. (C) The same field of view imaged as a medium magnification EM map. (D) Examples of individual cells with all possible barcodes, each column corresponds to one fluorescent channel, the rightmost column shows the same cell on the EM montage. Scale bars: 10 μm in B and C, 2 μm in D.

We call our new methodology MultiCLEM (for multiplexed CLEM). This methodology opens the door to new possibilities in cell biology where ultrastructural questions can be asked at a throughput previously only available for gross morphological changes in the cell.

## Results

### Fluorescent labeling of the cell wall enables molecular barcoding

Sample barcoding is a common parallelization approach in biology (Knapp et al., 2012; Krutzik and Nolan, 2006; Smith et al., 2009), and combinatorial fluorescent labeling is a powerful way to discriminate objects within heterogeneous samples (Livet et al., 2007; Valm et al., 2012). We therefore decided to harness these approaches to multiplex biological sample preparation for EM. We selected fluorophores that are very bright, retain their fluorescent signal after EM sample preparation, stain the same compartment consistently in different fluorescent channels, and can be easily delivered to the stained compartments in a combinatorial way. Fluorescent conjugates of Concanavalin A (Con A) that stain the yeast cell wall fulfilled these requirements. Con A conjugates in multiple colors are commercially available, and additional dyes can be easily attached using various protocols (Toseland, 2013). We selected five Con A conjugates (colors) that can be resolved on many conventional wide-field microscopes: Alexa350 (blue), Alexa488 (green), tetramethylrhodamine (TRITC) (orange), Alexa647 (far red), and Cy7 (near infrared).

To obtain specific barcodes, the selected Con A conjugates are mixed with each other in all possible combinations. Each particular conjugate (color) can be either present or absent in the mix (barcode). The total number of conjugates used depends on number of available fluorescent channels in the LM. More conjugates allow for more barcodes. (Fig. 1A). To increase the accuracy of barcode determination we did not use the possibility to mix Con A conjugates in different ratios and excluded the combination with no colors present to avoid false-negatives. This gives maximum seven (2^3^-1), 15 (2^4^-1), and 31 (2^5^-1) barcodes for three, four, and five Con A dyes respectively.

We optimized dye mix preparation and staining conditions to achieve the bright and uniform cell wall staining necessary for automated image processing. This resulted in an easy-to-use barcoding protocol (See Materials and Methods and Supplementary Data 1 for complete protocol) that could be followed by sample fixation, resin embedding, and LM imaging using a protocol that optimizes both preservation of fluorescent signals and cellular ultrastructures (Kukulski et al., 2012).

To illustrate the barcoding principle, we prepared a set of yeast cultures, using three Con A conjugates (Alexa 350, Alexa 488, and TRITC) combined in seven possible ways as described above. These cultures were mixed together in a single sample, which was subject to EM sample preparation and LM (Fig. 1B-D). When the three fluorescent channels are displayed as a pseudocolor overlay of red, green, and blue it is easy for the human eye to distinguish the seven possible combinations of three primary colors (red, green, blue, cyan, magenta, yellow, white), and thereby identify the source culture for each cell imaged by EM (Fig. 1D).

Manual analysis of images for sample identification is time consuming and restricts expansion of this methodology to high-throughput applications. We therefore developed a workflow for automation of barcode reading, correlation, and targeting of high-resolution acquisition (Fig. S1, S2). The workflow is organized in a *Matlab* graphic user interface (GUI) and allows correlation of LM data to medium magnification EM data for determination of cell barcodes with a possibility of selecting individual cells for subsequent automated high-magnification EM imaging (for detailed workflow description see Materials and Methods; the code and associated documentation is available at https://github.com/ybyk/muclem and https://www2.mrc-lmb.cam.ac.uk/groups/briggs/resources). Together, our barcoding approach and the software associated with it allow automated image acquisition of hundreds of cells from multiple samples for image analysis and phenotypic profiling.

### MultiCLEM allows quantitative ultrastructural phenotyping

The advantage of our proposed approach is that it allows to study ultrastructural phenotypes and to quantify parameters inaccessible to LM, while reducing time spent on sample preparation and data acquisition. To assess the developed MultiCLEM pipeline we chose to compare how the fine ultrastructure of membranous organelles is altered in different genetic backgrounds subjected to stress conditions.

We selected seven yeast strains: a reference lab strain S90 (Steinmetz et al., 2002), two different wine strains PRICVV50 (Novo et al., 2009) and SFB2 (Padilla et al., 2016), and four deletion mutants with defects in stress response and endomembrane system organization *Δhog1, Δtcb1, Δvps35, Δlpl1*. All strains were grown in parallel and then either subjected to hyperosmotic shock (1 M sorbitol for 45 minutes) or not. The 14 resulting yeast cultures underwent MultiCLEM (Fig. 2A) using combinations of four fluorophores.

**Fig. 2.**
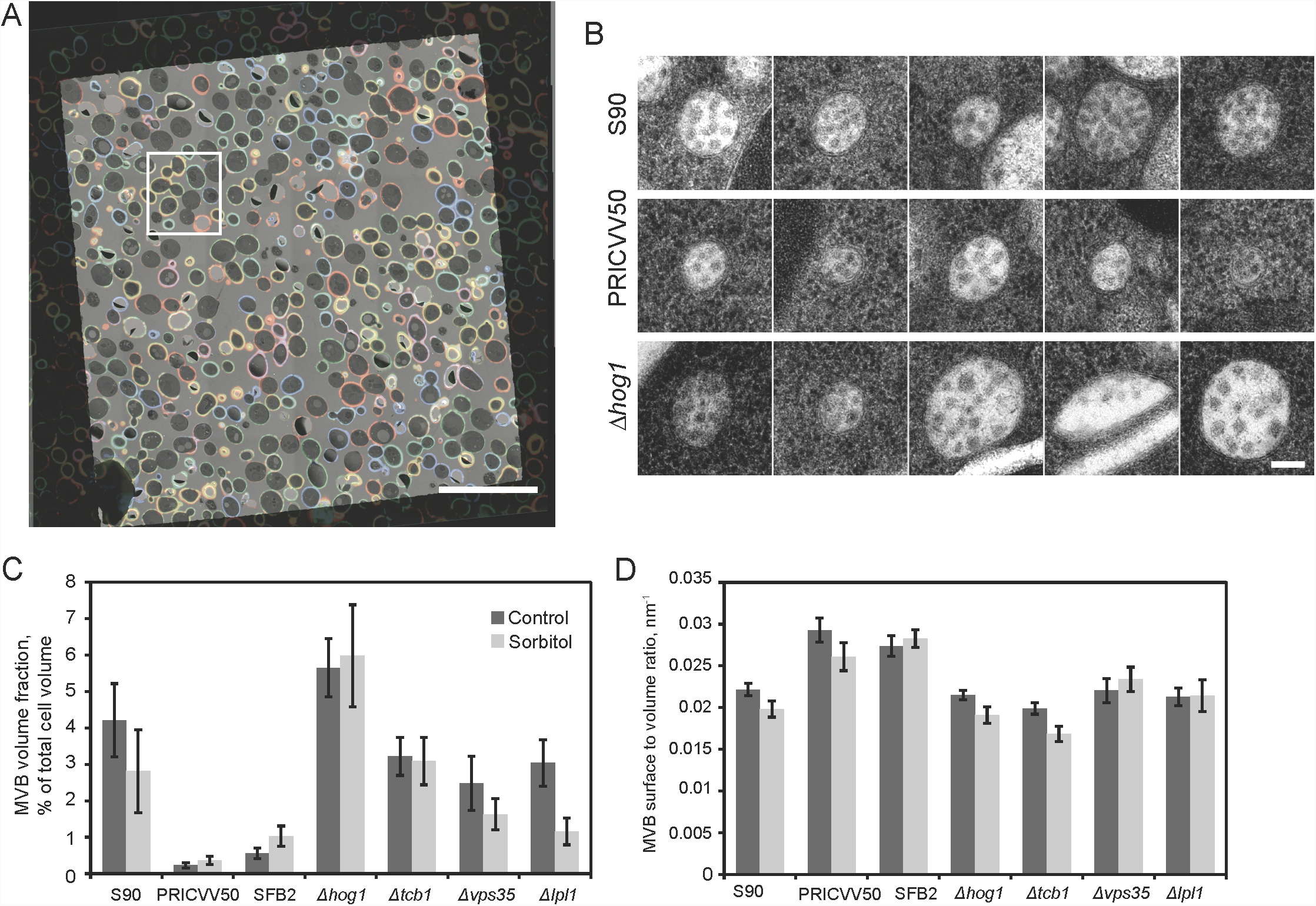
Ultrastructural phenotyping of yeast with different genetic backgrounds subjected to osmotic shock. (A) Overlay of pseudocolor composite fluorescent barcode image and medium magnification EM map of one grid square of the sample. The region in white rectangle is shown in detail on Fig S2. (B) Representative MVB cross-sections from the lab strain S90, wine isolate PRICVV50, and *Δhog1* strain. (C) MVB volume fractions in total cell volume in different strains and conditions, error bars show standard error of mean calculated as described in Materials and Methods. (D) MVB outer membrane surface to volume ratio in different strains and conditions, error bars show standard error of the mean. Scale bars: 20 μm in A, 100 nm in B.

All 14 barcodes were successfully recovered by the computational pipeline (Fig. S3). We first assessed the reliability with which cells were assigned to the correct strain. For 1000 cells we performed a careful visual assessment of the fluorescence signals to verify the barcode, generating a “gold standard” dataset which we compared to automated assignment: 87 % of cells were assigned to the same class visually and automatically, while 4 % of cells were incorrectly assigned by the automated procedure. 8 % of cells could not be classified visually due to the absence of cell wall staining or the signal obscured by aggregated Con A and thus automatic assignment was random. While such a low false positive and negative rate may be acceptable for high-throughput analysis if the sample size is big enough, we chose to create an easy tool for quality controlling and perfecting assignments for applications where higher accuracy is desired. We created an automated tool for examination of the dataset that enables, in a few hours, assessment of hundreds of cells and manual re-assignment or exclusion (the code and associated documentation is available at https://github.com/ybyk/muclem and https://www2.mrc-lmb.cam.ac.uk/groups/briggs/resources).

We then proceeded with high-magnification data collection. Previously-assigned cells could be identified for high-magnification EM imaging in a precise and error-free way using SerialEM version 3.7 (Mastronarde, 2005). Once set up, high-magnification imaging can run for a few days fully automatically. On different instruments high magnification data collection speed ranged from 30 to 70 cell cross-sections per hour at magnifications with pixel size ∼ 1 nm. The maximal number of cells per strain that can be imaged is obviously dependent on data collection speed, length of data collection, and number of strains within the multiplexed sample, but more than 100 cell cross-sections per strain can be attained routinely on standard electron microscopy set-ups.

For the experiment described above we acquired images of 1748 cell sections from which we generated galleries of around 100 high-resolution micrographs per cell strain and experimental condition (Fig. S4). The overall time needed to complete the described experiment was similar to that required for a single EM or CLEM experiment (Kukulski et al., 2012). While a small amount of additional work was required for fluorescence imaging, assigning strains and performing correlations, the increase in throughput of the most laborious steps was dramatic – fourteen samples were sectioned simultaneously, and inserted into the microscope in one step and imaged in an automated manner in only two stages: a short session to acquire medium magnification maps (2-3 h) and a long session for high-magnification imaging (around 24 h). Since it is feasible to perform 2-5 such EM experiments in parallel during a 2-3 week period, this makes it feasible to now study tens or even hundreds of strains where before only a few strains could be analyzed in a similar time period.

After confirming the effectiveness of our protocol, we examined the resulting high-resolution dataset. Yeast ultrastructure and preservation was similar to previously published data for healthy yeast cells in exponential growth phase that underwent high-pressure freezing and freeze-substitution (McDonald, 2007), meaning that yeast ultrastructure and preservation was not seriously affected by our barcoding protocol. While all EM embedding protocols may modulate ultrastructure to some extent, our approach optimizes the ability to compare strains because the control strain is processed within the same multiplexed sample.

By visual inspection, most organelles had a similar ultrastructural phenotype in all strains. However, striking differences were observed in the morphologies of mitochondria and multivesicular bodies (MVBs) when comparing to the reference strain (Fig. 2B, Fig. S5A). Mitochondria in the wine yeast strain SFB2 were swollen and lacked electron density in the matrix in both control and osmotic stress conditions (Fig. S5A). We focused our attention, however, on the structural changes observed in MVBs.

We decided to utilize the possibility presented by EM, allowing quantitative measurements of cellular features at the resolutions inaccessible to LM. Inspired by MVB morphology differences observed during visual examination of the dataset we localized and measured 510 MVB cross-sections in 1748 imaged cell cross-sections, and estimated an average volume fraction and surface to volume ratio of these organelles (Fig. S5B,C, see Materials and Methods for details). We found dramatic variation in the fraction of cellular volume taken up by MVBs (MVB volume fraction) (Fig. 2C) while the surface to volume ratios were relatively uniform (Fig. 2D).

Little is known about the MVB size, shape and abundance regulation in yeast, despite the fact that the basic mechanisms of MVB biogenesis were uncovered in this model organism (Katzmann et al., 2001; Hanson and Cashikar, 2012; Nickerson et al., 2010; Arlt et al., 2015). Usually, with increase of cell size, including the transition from haploid to diploid, the fraction of cellular volume taken up by different organelles tends to stay constant or show a slight increase (Chan and Marshall, 2010). In contrast, we observe dramatically reduced MVB volume fraction in diploid wine yeast strains with larger cells (Fig. 2C, S5B). This must reflect the different genetic background of the wine yeast strains: genome sequencing and phenotyping reported that strains of such origin differ significantly in their physiology from beer and laboratory yeast strains which have been cultured in rich media for many generations and, for example, have lost many stress-response capabilities (Gallone et al., 2016). Our method can be useful for subsequent study of this topic, and for exploration of other organelles similar to MVBs, whose volumes and sizes cannot be precisely measured using LM.

### Fluorescent barcoding CLEM as a part of two-step screening strategy

The ultimate goal of MultiCLEM in yeast is to be used as a screening platform for cell traits not screenable with the standard techniques based on LM. At present, our set up cannot support the screening of an entire yeast deletion library although future developments (see discussion below) could in principle bring it to a scale compatible with whole-genome screens. Hence a current, valuable, utilization of MultiCLEM could be in secondary screens where the primary LM screen narrows down the number of strains and then a secondary screen to verify hits can be performed by MultiCLEM. As proof of principle of this perspective use, we performed a primary LM screen followed by a secondary MultiCLEM screen on a cellular phenotype not easily discernable by LM – mitochondrial ultrastructure.

The mechanisms underlying mitochondrial fission and fusion have been well worked out in yeast and are highly regulated and interrelated processes (Friedman and Nunnari, 2014). Surprisingly, however, it is still not clear how mitochondrial width and branching are regulated or determined. Moreover, even in the complete absence of both fission and fusion machineries, mitochondria still display differing morphologies, branching and dynamics suggesting that additional mitochondrial shaping proteins exist. To uncover such proteins we performed a two-stage screen consisting of a higher-throughput live-cell fluorescent microscopy stage to select initial hits followed by a lower-throughput high-resolution characterization of the ultrastructure of these hits using MultiCLEM.

In the first stage we imaged mitochondria labeled with endogenous Tom20 fused with GFP in over-expression strains of 99 mitochondrial proteins (enriched for outer membrane proteins) on the background of a deletion of Dnm1 – the yeast dynamin-like protein in whose absence the major fission events cannot take place (Fig. 3A). *Δdnm1* mutant cells show dense, clumped mitochondrial networks by LM, while wild type (WT) cells show individual tubules connected into a loose network mostly positioned in the cell periphery (Fig. 3B). We expected that overexpression of membrane deforming proteins could suppress the defects of losing Dnm1 or alter mitochondrial morphology in some other way.

**Fig. 3.**
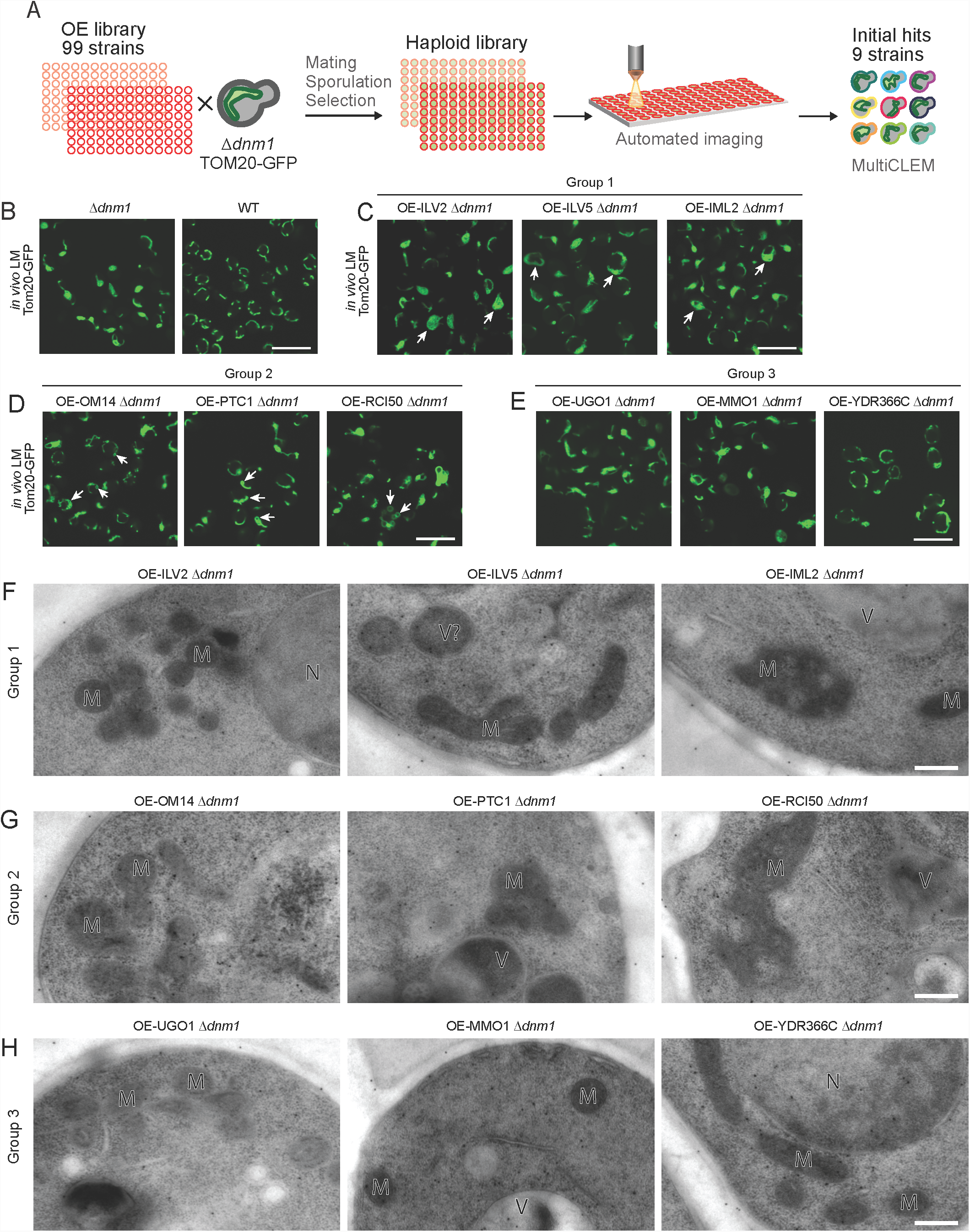
Identification of new mitochondrial shaping proteins using two-step screening. (A) The screen setup: Using automated mating, sporulation and haploid selection approaches a tailor made library of 99 strains was created such that all strains had a background of both Δdnm1 and TOM20-GFP (as a mitochondrial marker) as well as an overexpressed (OE) allele of one mitochondrial associated protein (enriched for outer mitochondrial membrane). Strains were then imaged using live LM. Nine strains were selected for EM analysis. (B)-(E) Mitochondrial morphology visualized by LM in living cells expressing Tom20-GFP. (B) The overexpression background strain Δdnm1 TOM20-GFP compared to the WT TOM20-GFP strain. (C) Group One strains in which a subset of cells have expanded mitochondrial networks (arrows). (D) Group Two strains in which circular, GFP-positive compartments are observed (arrows). (E) Group Three strains in which a subset of cells have WT mitochondrial morphology indicating a partial Δdnm1 rescue phenotype. (F)-(H) Characterization of the mitochondrial ultrastructure using MultiCLEM. (F) Group One strains. (G) Group Two strains. (H) Group Three strains. Organelles marked in EM images: M – mitochondria, V – vacuole, N– nucleus. Scale bars are 10 µm for LM images, and 200 nm for EM images.

After visual assessment of the LM images of the 99 strains in the overexpression mini-library, we selected nine strains with mitochondrial morphologies distinct from the *Δdnm1* strain. At LM resolution they can be divided into three groups. Group One (overexpressing Ilv2, Ilv5, and Iml2) was characterized by the expansion of the clumped mitochondrial networks manifested by the appearance of a small number of cells with hyper-proliferated mitochondria that occupied most of the confocal cross-section area (Fig. 3C). Group Two (overexpressing Om14, Ptc1 and Rci50) possessed small, circular Tom20-GFP positive structures with diameters up to 500 nm in addition to mitochondria with *Δdnm1*-like morphology (Fig. 3D). Group Three (overexpressing Ugo1, Mmo1, and Ydr366c) showed a partial ‘rescue’ phenotype in which some cells had close-to-normal, extended mitochondria (Fig. 3E). The strain overexpressing Ydr366c had the most pronounced rescue phenotype with many mitochondria resembling those in WT cells.

In the second stage of the experiment we used MultiCLEM for ultrastructural characterization of the previously selected nine strains (Fig. 3F-H, Fig. S6). We prepared ultrathin sections and collected high-resolution 2D EM micrographs of ∼ 60 cell cross-sections for each of the nine strains. Using the data we could assess the diameters of mitochondrial tubules, their electron density and the presence of other unusual structures in mitochondria or in the cell in general. We aimed to determine whether overexpression of the selected proteins caused side effects resulting in altered mitochondrial ultrastructure or other abnormalities, which might suggest that the phenotype observed by the LM screen is unspecific.

We did not find any dramatic mitochondrial ultrastructure alterations such as those observed in wine strain SFB2 (Fig. S5A). Most individual mitochondria in all strains had an ultrastructure similar to that in WT cells (Fig. S7): circular or extended structures 200-400 nm in diameter with electron density higher than that of the cytoplasm. In *Δdnm1* cells, mitochondria are often found in large clumps.

Group One strains had an ultrastructural phenotype similar to the WT (Fig. 3F, Fig. S7). In the Group Two strains, we did not encounter any mitochondria that might correspond to the circular structures visible by LM: the mitochondria had neither a very large diameter comparable to the size of these structures, nor did they form any circular clumps. However, in a small number of cross sections we observed compartments more similar to vacuoles by texture and bounding membrane ultrastructure, that were of a similar size to the structures observed by LM (Fig. 3G, Fig. S8). Immunogold labeling showed that these structures do indeed contain Tom20-GFP (see next section) suggesting that the phenotype we saw by LM occurred as a result of increased mitophagy. In Group Three, which partially rescued the *Δdnm1* phenotype by LM, some cells overexpressing Ugo1 had mitochondria with lower electron density and also showed vacuole-like structures of irregular shape that were labeled by anti-GFP antibody (see next section). The strains overexpressing Mmo1 and Ydr366c had mitochondrial sizes, distribution and electron density, as well as overall cell morphology similar to that of WT (Fig. 3H), suggesting that overexpression of these two proteins rescues the *Δdnm1* mitochondrial phenotype without side effects affecting cellular ultrastructure.

To summarize, as result of this screen in which we combined conventional LM screening and a secondary ultrastructure characterization using MultiCLEM, we identified Mmo1 and Ydr366c as potential factors that might play an additional role in establishing mitochondrial morphology. Both of them are small proteins that localize to mitochondria when tagged with GFP (Yofe et al., 2016), have an unknown function, and have predicted trans-membrane helices.

### Multiplexed barcoding combined with GFP and immunogold labeling for organelle identification

Both mitochondria and MVBs have distinct features making them clearly identifiable by EM, however many organelles such as peroxisomes or small vesicles are not easily identified either by eye or computationally. Hence, to enable our method to be quantitative for a large number of traits we sought additional means to accurately assign identity to a variety of cellular structures. Currently two ways exist to do this – immunogold labeling and correlative microscopy – and we chose to explore both approaches.

#### Immunogold labeling

We optimized an immunostaining protocol to visualize any protein fused to GFP with anti-GFP antibodies on our multiplexed samples. We then used this protocol to perform immunogold labelling on the set of *Δdnm1* strains expressing Tom20-GFP and overexpressing the nine mitochondrial outer membrane proteins described in the previous section. This was necessary to analyze the strains overexpressing Rci50, Om14, Ptc1, and Ugo1 that had structures resembling small vacuoles in addition to mitochondria with normal morphology (Fig. 3G, H, Fig. S8). Immunogold labeling confirmed that these structures indeed contained GFP. The specificity of immunolabeling was confirmed by quantifying the number of gold beads localized to mitochondria and vacuoles in the strains where these organelles can be unambiguously identified visually. In 60 quantified cell cross-sections 74% of beads localized to mitochondria; the other 24% of beads were preferentially localized to the vacuole and cell wall. This is much higher than the 5-15% of the cross-section area that is occupied by each of these organelles (Rafelski et al., 2012; Chan and Marshall, 2014). Hence, immunostaining can be used with multiCLEM to help to define cellular structures.

#### Correlative microscopy

To perform in-resin CLEM, we focused on peroxisomes. Peroxisomes are small organelles with diameters smaller than the resolution of LM. We expressed a peroxisomal targeted fluorophore (Grx1-GFP-PTS1) in 14 yeast strains each carrying a deletion of one peroxisomal protein as well as in a control strain (see Table S1 for the full list). The strains were grown on glucose-containing media (where peroxisomes are fewer and smaller and hence harder to visualize, as a proof of principle for the effectiveness of the CLEM approach), barcoded using four Con A conjugates, and processed for EM. We successfully identified all 15 strains in fluorescent micrographs, and punctate GFP signals were visible in all strains except the *Δpex22* strain which did not correctly target this reporter to peroxisomes under these growth conditions. After correlation and analysis of 518 cells at medium and high resolution we confidently identified 46 structures as peroxisomes in 11 strains out of 14 that showed punctuate signals in LM (Fig. 4). To speed up the analysis we did not add fluorescent fiducial markers for correlation. That resulted in low correlation precision, and we were unable to confidently correlate some signals coming from cells belonging to the three remaining strains. The peroxisome cross-section diameters varied from 100 to 400 nm (Fig. 4B) and some of the smaller peroxisomes showed increased electron density of the contents (Fig. 4A). Peroxisomes tended to be larger in the strain lacking Ant1 (a peroxisomal ATP transporter) demonstrating the power of this approach to differentiate ultrastructural details that can’t be uncovered using conventional LM screening and image analysis.

**Fig. 4.**
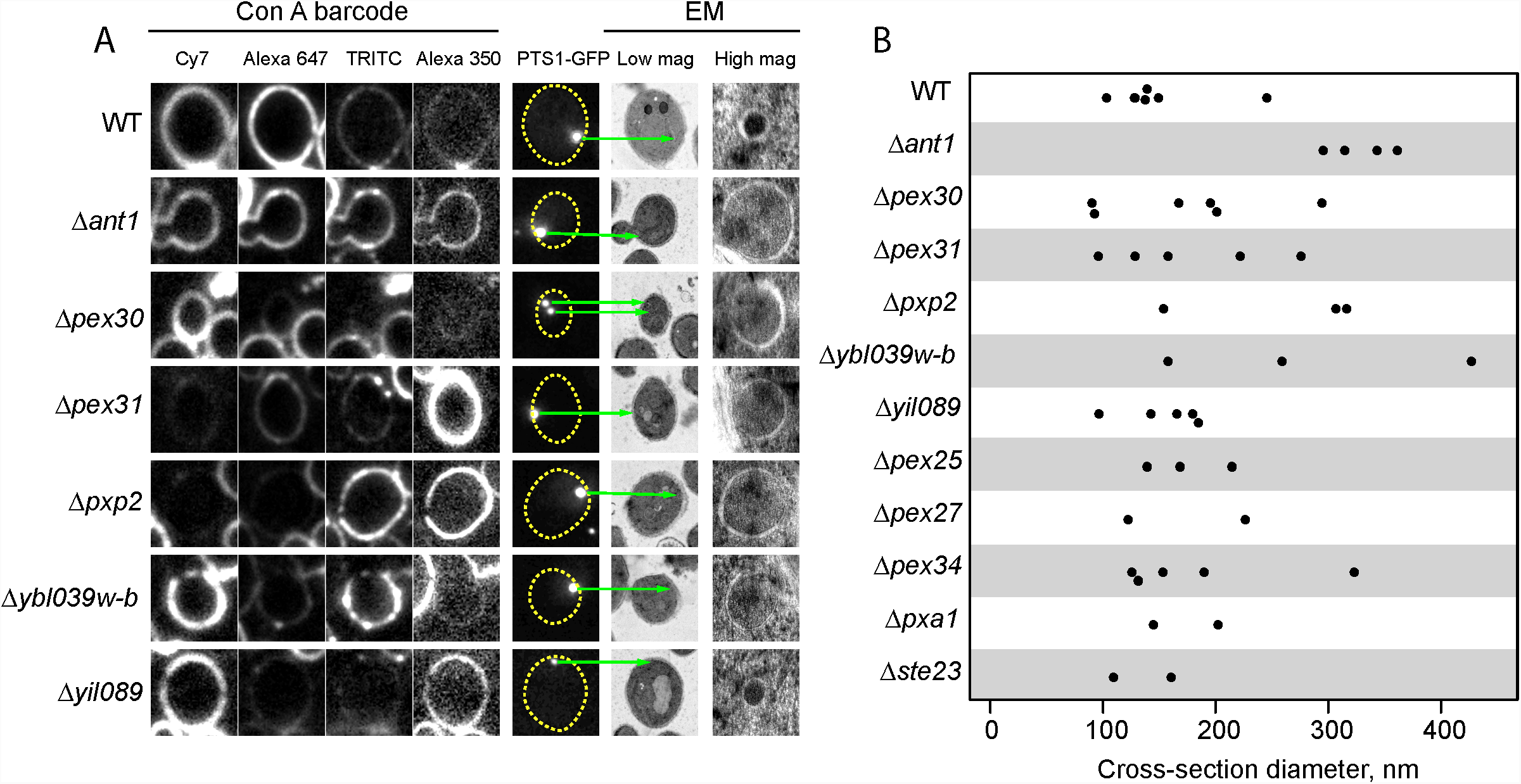
Identification of peroxisomes in different deletion strains using CLEM. (A) An example set of cell cross-sections where peroxisomes could be identified in the high-resolution data: first four columns show images taken in each fluorescent channel from which the barcode is read; fifth column shows the GFP signal; sixth column shows EM images of the corresponding cell; seventh column shows high-magnification (mag) EM images at the position of the GFP puncta. (B) Peroxisome cross-section diameters measured for all identified peroxisomes in different strains. Size of all LM and low magnification LM panels is 5 μ m, high mag EM panel is 400 nm.

## Discussion

In this work we describe a new approach allowing systematic, parallel, high-throughput screening for traits observable by EM. We demonstrated that parallel processing of up to 15 barcoded strains within a single EM sample can be performed on a timescale similar to that of a typical EM or in-resin CLEM experiment. A typical automated freeze-substitution processor can produce 10-20 resin blocks within one week and they can be sectioned within few days. Given two blocks per sample, five to ten multiplexed samples can be processed and sectioned in parallel allowing 70-140 strains to be assessed using a single freeze-substitution run. Hundreds of EM images of each strain could then be collected in an automated manner within an extended EM imaging session. Such an experiment using conventional approaches would take months of consecutive freeze-substitutions and many days of laborious sectioning and EM imaging.

Because multiple strains are treated in one sample, all cells belonging to the different strains have been prepared for EM and imaged in parallel. This eliminates any variation that arises from differences in vitrification during high-pressure freezing, sample shrinkage during freeze-substitution, compression during sectioning, contrast and brightness during post-staining and imaging. If necessary a control sample can be included in every multiplexed experiment for direct comparison hence our method not only accelerates data acquisition but also increases its accuracy. The parallel nature of the experiment means that quantitative comparative analysis of immunogold-labeled sections can be performed. It also facilitates automated analysis of the images allowing direct quantitative comparison of strains using image processing algorithms or neural networks.

In our protocol we used five spectrally non-overlapping fluorophores (four for barcoding and one for GFP). In principle, all five can be used for barcoding if cells contain no additional fluorescent tags. Each fluorophore had two possible staining levels (stained or not stained) giving a maximum number of 2^5^-1=31 combinations (the combination with no staining for all fluorophores is excluded for accuracy purposes). It is reasonable to prepare five samples in parallel which would provide for up to 155 strains. A further increase in the number of multiplexed experiments could be achieved by several means. More fluorophores with overlapping spectra could be used, and linear unmixing applied. Different staining levels could be used for each fluorophore similar to existing fluorescent barcoding techniques like BrainBow and cell fluorescent barcoding for cell sorting (Weissman and Pan, 2015; Krutzik and Nolan, 2006). Internal organelles could be stained in the same combinatorial manner, in addition to the cell wall, providing the opportunity for hundreds of samples to be barcoded at once. If that were to be developed, the available electron microscope time to acquire high resolution images of all such strains in a sample would quickly become limiting. In the future, such limitations could be overcome by increasing data collection speed using multi-beam EM (Lena Eberle et al., 2015) or by precise targeting of data collection using CLEM or immunogold labeling so that only regions of interest are imaged.

Our protocol is currently adapted for budding yeast but it can also be used for other species that can be stained with Con A, including some mammalian cell lines. Similar protocols can be developed for bacterial or mammalian cells using antibodies labeled with various fluorophores, chemical dyes or genetically encoded fluorescent proteins.

The approach that we have presented here allows both highly multiplexed experiments, and parallel sample preparation. It therefore fulfills the requirements for quantitative screening of EM samples. We believe that this approach will lay the foundation for expanding systematic screening and high-throughput imaging approaches to the ultrastructural level using CLEM.

## Acknowledgements

We are grateful to Markus Mund and Albert Mas for yeast strains and to Nadav Shai, Maria Bohnert and Michal Eisenberg-Bord for sharing unpublished reagents. We thank Morgane Wartel, Lisa Maier, Einat Zalckvar, Nassos Typas and Anne-Claude Gavin for advice and discussions. We are grateful to Michael Knop for discussions and comments on the manuscript. This study was technically supported by the EMBL Advanced Light Microscopy Facility, EMBL Electron Microscopy Core Facility, Weizmann Institute of Science Electron Microscopy Facility, and MRC LMB Light Microscopy Facility. We thank Martin Schorb, Yannick Schwab and Rachel Mellwig for their help and advice on setting up the MultiCLEM method at EMBL. This work was supported by the Deutsche Forschungsgemeinschaft within SFB1129 Z2 to JAGB, the European Molecular Biology laboratory to JAGB, the Medical Research Council MC_UP_1201/16 to JAGB, and German Ministry of Education and Research (BMBF, grant number 031A605) to KRP. The Schuldiner lab is supported by an ERC CoG 646604 Peroxisystem, the Deutsche Forschungsgemeinschaft within SFB 1190, and a DIP collaborative grant. NG was supported by the EMBL interdisciplinary postdoctoral program. MS is an incumbent of the Dr. Gilbert Omenn and Martha Darling Professorial Chair in Molecular Genetics.

## Materials and Methods

### Yeast strains

The yeast strains used in this study are listed in Table S1.

The library for the peroxisomal morphology screen was prepared by mating a roGFP-PTS1 plasmid containing query strain (constructed on the basis of a synthetic genetic array (SGA) compatible strain, YMS721 (Papić et al., 2013)) with a collection of ∼15 strains in which peroxisomal genes were deleted using a KanMx KO cassette (Goldstein and McCusker, 1999). Automated sporulation and selection of haploids was performed using the SGA method (Cohen and Schuldiner, 2011; Tong and Boone, 2006) in high-density format using a RoToR bench top colony arrayer (Singer Instruments).

A similar SGA approach was used to create the library for the mitochondrial morphology screen by mating a *TOM20-GFP, Δdnm1* expressing query strain to a collection of ∼100 strains in which genes for mitochondrial outer membrane proteins, and a number of other mitochondria-related proteins, have been modified to be driven by a *TET-OFF* promoter.

### Cell growth

For protocol development, cells were grown in YPAD medium or synthetic complete medium without tryptophan (SC-Trp). For parallel growths of more than 7 strains we used 24-well plates. To ensure equal gas exchange rate in all wells, the plate was sealed with a gas-permeable membrane (Sigma-Aldrich) and placed on a plate shaker set to 600 rpm to achieve proper mixing of the suspension. The shaker was in turn placed in a conventional incubator at 30°C. Most experimental cultures were inoculated from colonies on agar plates, grown overnight and in the morning diluted to OD_600_ 0.1-0.2. Diluted cultures were grown to OD_600_ 0.5-0.8 for assaying.

For MVBs comparison, strains were grown in parallel and then either subjected to hyperosmotic shock (1 M sorbitol for 45 minutes) or not.

For the peroxisome morphology screen the strains were grown in SD-MSG supplied with G418.

For the mitochondrial outer membrane primary fluorescent microscopy screen, the strains were transferred from agar plates into 384-well polystyrene plates for growth in liquid media using the RoToR arrayer robot. Liquid cultures were sealed with a gas-permeable membrane and grown in a shaking incubator (Heidolph Tiramax1000, inkubator1000), overnight at 30°C in YPAD medium containing Hygromycin (Hygro), Neurseothricin (NAT) and G418. To drive overexpression of the TET-off promoter no tetracycline was added to the medium. The strains were diluted to OD_600_ of ∼0.2 into plates containing YPAD medium and incubated for 4 hours at 30°C. The cultures in the plates were then transferred into glass-bottom 384-well microscope plates (Matrical Bioscience) coated with Concanavalin A (Sigma-Aldrich). After 20 minutes, wells were washed twice with YPAD to remove non-adherent cells and to obtain a cell monolayer.

For the mitochondrial outer membrane secondary EM screen, the hits from the primary fluorescent microscopy screen were grown in a 96 well plate, sealed with a gas-permeable membrane, in a shaking incubator (Heidolph Tiramax1000, inkubator1000), overnight at 30°C in YPAD medium. The strains were diluted to an OD_600_ of ∼0.2 into 96-well 2ml tall-well plate containing YPAD medium, sealed with a gas-permeable membrane, and incubated for 4 hours at 30°C.

### Live imaging of yeast mitochondria

Strains were imaged using VisiScope Confocal Cell Explorer system, composed of a Zeiss Yokogawa spinning disk scanning unit (CSU-W1) coupled with an inverted Olympus microscope (IX83; ×60 oil objective; Excitation wavelength of 488 nm for GFP and 560 nm for mCherry). Images were taken by a connected PCO-Edge sCMOS camera controlled by VisView software.

### Production of fluorescently labelled Con A

For cell wall staining we purchased Con A conjugated with Alexa Fluor^®^ 350, Alexa Fluor^®^ 488, Tetramethylrhodamine (TRITC), and Alexa Fluor^®^ 647 from Life Technologies. Stock solutions with concentration 2.5 mg/ml were prepared in phosphate buffered saline (PBS) and stored at −20°C in small aliquots. Con A conjugated to Cy7 stock solution was prepared using the following protocol (Mund et al., 2014). Sulfo-Cy7 NHS ester (Lumiprobe) was diluted in dimethylsulfoxide (DMSO) to a concentration of 10 mM. Concanavalin A (type IV, Sigma-Aldrich) 2.5 mg/ml solution was prepared in 0.2 M NaHCO3 with pH 8.2. Dye and protein solutions were mixed 6:100 and incubated for 4 hours at room temperature. Conjugated Con A was separated from the reaction on a disposable Sephadex G-25 column and buffer exchanged to PBS. The stock solution was stored frozen in small aliquots.

### Barcoding

Con A stock solutions were diluted with PBS and mixed in different combinations to yield the final staining solutions (See Supplementary data 1 for details). Yeast strains grown to logarithmic phase were pelleted by centrifugation of the multi-well plate for 5 minutes at 1500 rcf. The growth media was removed and the pellets were resuspended in the staining solutions. The cells were incubated in the staining solutions with shaking at 30°C (the same as growth conditions) for 10 minutes. The cells were pelleted for 5 minutes at 1500 rcf and resuspended in YPD media. All contents of the multi-well plate were mixed together, thoroughly vortexed, and immediately processed for EM. A detailed protocol is provided in the Supplementary data 1.

### EM sample preparation

We followed a standard sample preparation protocol used for in-resin CLEM (Kukulski et al., 2012). Immediately after barcoding, the yeast biomass was collected using a Millipore filtering setup on a 0.45 μm nitrocellulose filter. The cell slurry was transferred to the 0.1 mm deep cavity of a 0.1/0.2 mm membrane carrier for an HPM010 high-pressure freezing machine or Leica ICE. The cavity was covered by the flat side of a 0.3 mm carrier, and the sandwich was inserted in the high-pressure freezing machine. Resin embedding was performed using a Leica AFS2 freeze-substitution machine equipped with a processing robot. Samples were embedded in Lowicryl HM20 resin using the freeze-substitution and embedding protocol optimized for in-resin CLEM (Kukulski et al., 2012). Dry acetone with 0.1 % uranyl acetate was used as the freeze-substitution medium (FS medium). The blocks were trimmed with a razor blade and 100 nm thick sections were produced using a Diatome 35° knife on a Leica Ultracut UCT or UC7 microtome. The sections were mounted on 200 mesh copper grids with continuous carbon support film (Electron Microscopy Sciences). Grids were imaged under the light microscope the same day.

### Fluorescence microscopy for CLEM

Fluorescence microscopy was performed using a protocol for in-resin CLEM imaging (Kukulski et al., 2012). The grid was sandwiched between two coverslips with a droplet of distilled water or PBS and mounted on the microscope stage using a holder. We used a Nikon TE2000 microscope for the peroxisome morphology screen, a Zeiss Cellobserver Z1 for mitochondria morphology experiment, and a Zeiss Cellobserver Z1 for all other experiments. Filter sets and other characteristics for each setup are outlined in Table S2. Usually, five to ten positions (grid squares) were imaged on each grid. Between those, exposure times and other conditions were kept the same. After imaging, coverslips were separated and the grid was carefully recovered and dried. If the imaging was performed in PBS the grid was washed in distilled water before drying.

### Immunogold labeling

Following the in-resin CLEM imaging, the grid was washed twice for 3 minutes in washing solution containing filtered PBS and 0.2% glycine (Sigma-Aldrich), blocked for 30 minutes in filtered blocking solution containing: 0.5% gelatin (EMS), 0.5% BSA (MP biomedical) and 0.2% glycine (Sigma-Aldrich), in PBS. Then samples were incubated for 2 hours with AB290-rabbit anti-GFP antibody (Abcam) diluted 1:50 in filtered blocking solution, washed five times for 2 minutes in washing solution, blocked for 5 minutes in filtered blocking solution, incubated for 45 minutes with goat anti-rabbit 15nm gold antibody (EMS) diluted 1:20 in filtered blocking solution, washed five times for 2 minutes in washing solution, washed three times for 2 minutes in filtered double distilled water (DDW) and then the grid was kept on a drop of DDW until post-staining.

### Electron microscopy

Prior to EM, grids were post-stained for 2 minutes in Reynolds lead citrate. For the mitochondria morphology screen, EM imaging was performed on a T12-Spirit Bio-Twin electron microscope (FEI) operating at 100 kV, equipped with an Eagle 2k × 2k detector (FEI). For all other experiments, EM imaging was performed on a TF30 electron microscope (FEI) operating at 120 kV, equipped with a Gatan OneView detector. SerialEM software was used for data collection (Mastronarde, 2005), and the detector was operated in the full frame mode (4096 × 4096 pixels in the TF30 and 2048 × 2048 pixels in the T12). Regions imaged using LM were localized and maps were produced at magnification 2000-4000X (“medium magnification” montages, correspond to pixel size 2.5-4 nm). Complete imaging of each region (containing one grid square) required a 5×5 to 13×13 montage. The resulting montages were saved as maps in the SerialEM Navigator file and used for identification of cells during image analysis and correlation. After processing, cell positions for high-resolution imaging were imported to the Navigator file and used to acquire a 2×2 montage of each cell with magnification 9400X (“high magnification” images, correspond to pixel size ∼ 1 nm). Additional practical instructions for setting up the EM imaging are provided in the software manual.

### Image analysis and correlation pipeline

Here we describe the general image and data processing workflow we used. For a complete description of the software refer to the manual provided with it. The code and documentation is available at https://github.com/ybyk/muclem and https://www2.mrc-lmb.cam.ac.uk/groups/briggs/resources.

#### Cell detection

We used medium magnification montages to obtain positions and outlines of cells. Montages were blended using *blendmont* software from the IMOD package (Kremer et al., 1996) to produce a single grid square overview. This overview was segmented using the pixel classification workflow in *ilastik* software (Sommer et al., 2011). *Ilastik* was used to reliably distinguish well-preserved cells from resin, holes in resin where cells had detached, and dark regions containing cell debris. The segmentation was loaded into *Matlab* (Mathworks) and all processing was performed using Image Processing Toolbox functions. Cell outlines were produced by smoothing the raw *ilastik* segmentation and applying watershed segmentation. Cells were automatically identified based on the object size and circularity. Outline and center coordinates for each cell were saved.

#### Fluorescence intensity measurement

The *Matlab* control point selection tool was used to correlate LM and EM images and calculate the transformation between them. Affine transformation using 5-10 registration points was sufficient to overlay EM-derived outlines with fluorescent data and collect all signal for most cells. Cell outlines determined from the EM montages were converted to masks by a morphological dilation procedure. The intersections of masks with masks of neighboring cells and holes were subtracted from each mask to avoid measurement bias. The resulting masks were then applied to the LM images, and the fluorescent signal was measured within the masks.

#### Barcode determination

For each cell, median intensity was measured in each fluorescent channel within the mask. These values were normalized in two steps. First, to bring all channels to similar ranges and intensity distributions, values for each cell in each channel were normalized by subtracting the minimum observed value in this channel in all cells and dividing all values by interquartile range:

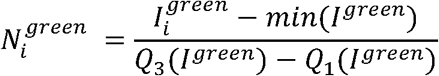

Where 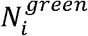 is the normalized intensity of the i-th cell in the green channel (as an example), 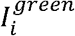 is the raw intensity of this cell measured using the mask, 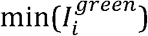 is the minimal raw intensity observed in the green channel for all cells, 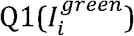 and 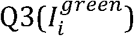are the lower and upper quartiles of intensities observed in the green channel for all cells. Interquartile range was used instead of minimum-maximum range because it better characterized the distribution shape and did not depend on outliers.

Since all cells displayed a variable total amount of labeling (but highly correlated intensity between channels), we normalized intensities the second time for each cell between channels by dividing all values for each cell by the value of the channel with maximal intensity:

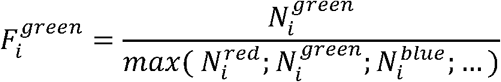

Where 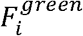 is final normalized intensity of i-th cell in green channel and 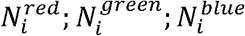 … are normalized intensities in red, green, blue, and other channels calculated as described above. After the second normalization, k-means clustering was performed on the final normalized intensities with the number of clusters corresponding to the expected number of color combinations.

The coordinates of cells selected for high-resolution acquisition were imported to the SerialEM Navigator file and high-resolution EM images were collected of these cells. High-resolution images were blended into montages using blendmont, the cells were sorted according to their staining patterns. All data analysis was performed in *Matlab*.

### Multivesicular body morphometry

Each cell micrograph was examined in IMOD without knowledge of its barcode. If an MVB was present on the cross section, it was approximated by eye as an ellipse and its major axis and minor axis perpendicular to it (as the widest place in the direction perpendicular to the major axis) were identified – D_1_ and D_2_ respectively (Fig. S5C). Area (*A*) and circumference (*C*) of each MVB cross section were calculated according to the following formulas:

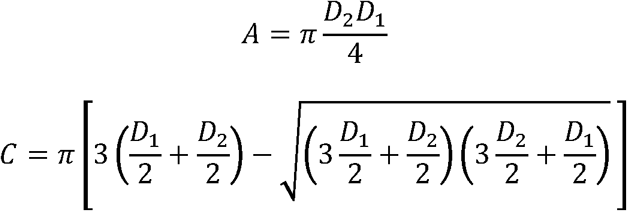

Areas of cell cross sections were determined from medium magnification montage segmented using *ilastik* (see above). The volume ratio of MVBs in total cell volume (*V*_*r*_) was determined using a well-known stereology relationship between the cross-section area occupied by the studied compartment and the total cell cross-section area, using the areas measured above instead of using traditional stereology point counts (Howard and Reed, 2005):

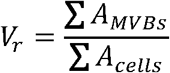

The standard error of this measurement was estimated according to the method described in Howard and Reed, 2005:

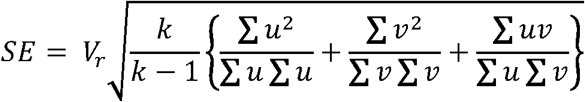

Where *k* is a total number of cross-sections, and *u* and *v* are vectors with values of cell cross-section areas and MVB cross-section areas for each cell, respectively. Each summation is over 1 to *k*. MVB surface to volume area was estimated as the relation of the sum of MVB cross-section areas to the sum of circumferences. Data processing and plotting were performed in *Matlab, Libre Office Calc*, and *R* (R Core Team, 2013).

---

**Fig. S1.**
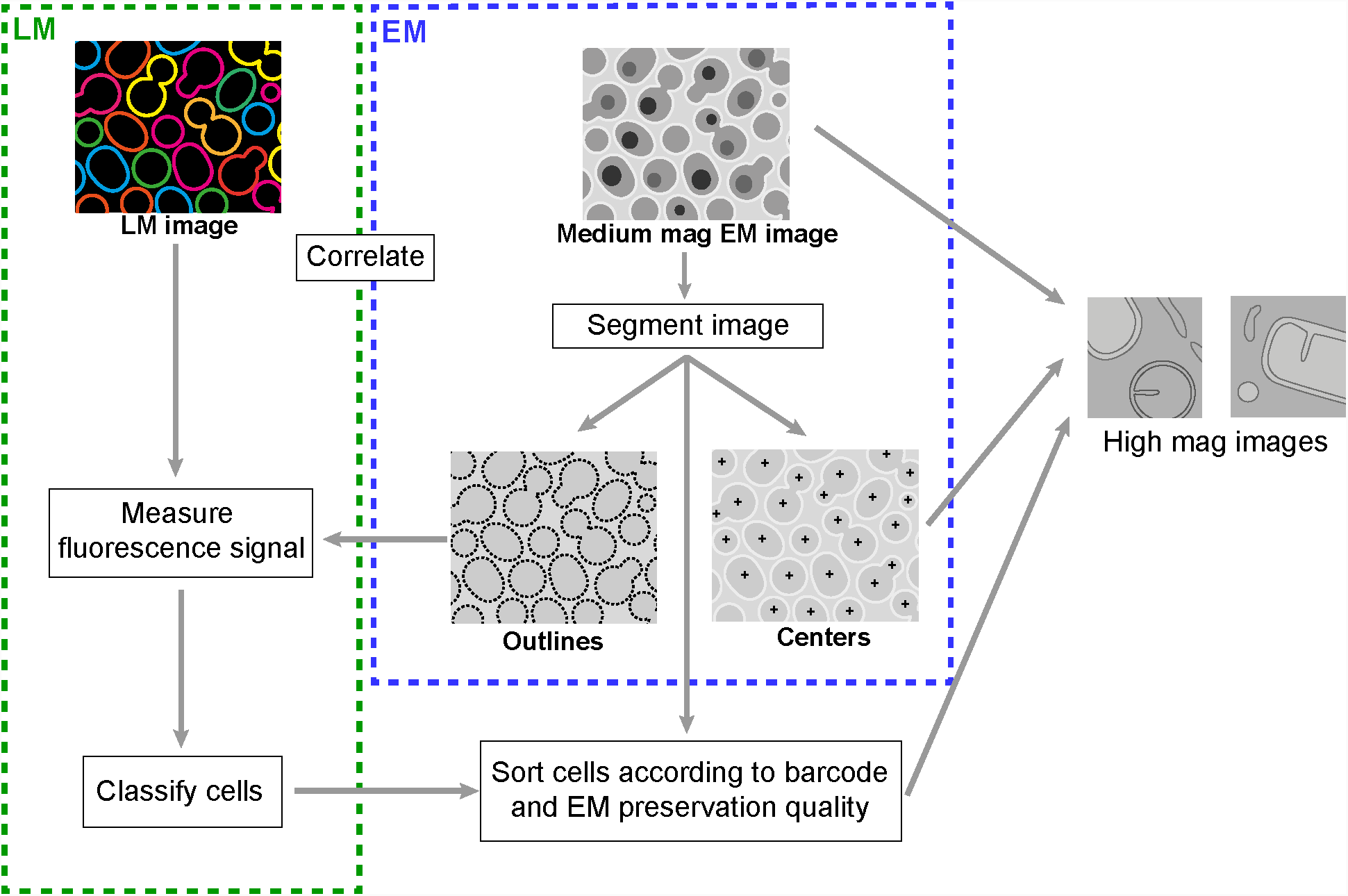
Image and data processing pipeline. Medium magnification EM images are segmented to derive cell centroids and outlines. Correlation of EM and LM data is performed, after which cell outlines determined from the EM data are used to create masks to measure fluorescence intensity. Cells are classified by fluorescence intensities and barcodes are determined. Cells with unknown barcode or poor preservation are excluded during a quality control step. Centroids of selected cells are imported back to the electron microscope for high-resolution imaging.

**Fig. S2.**
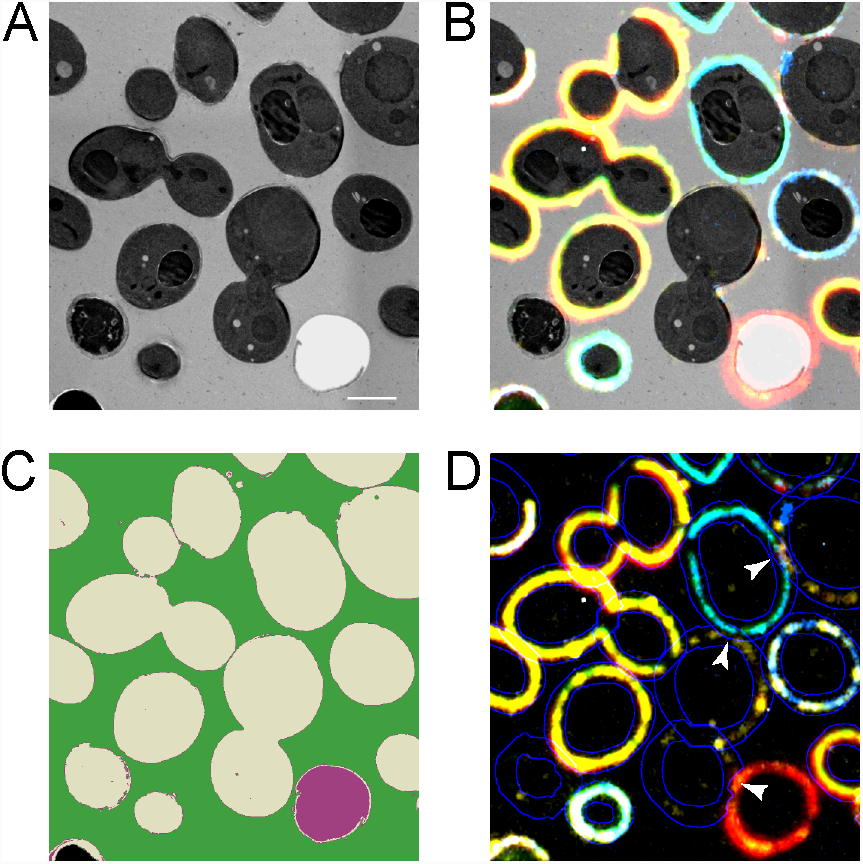
EM image segmentation and cell wall detection. (A) Close up view of part of the EM image shown in Figure 2A. (B) The same region as in A, overlaid with the fluorescent signal. (C) Ilastik segmentation of the same EM image. Cells suitable for imaging are shown in gray, resin in green, holes in the resin are in purple, electron-dense regions are in black. (D) Cell wall outlines determined from EM image segmentation and used to measure fluorescent intensities are shown in blue and overlaid with the composite fluorescent image, white arrow heads show some of the regions excluded from masks due to intersections with neighboring cell walls.

**Fig. S3.**
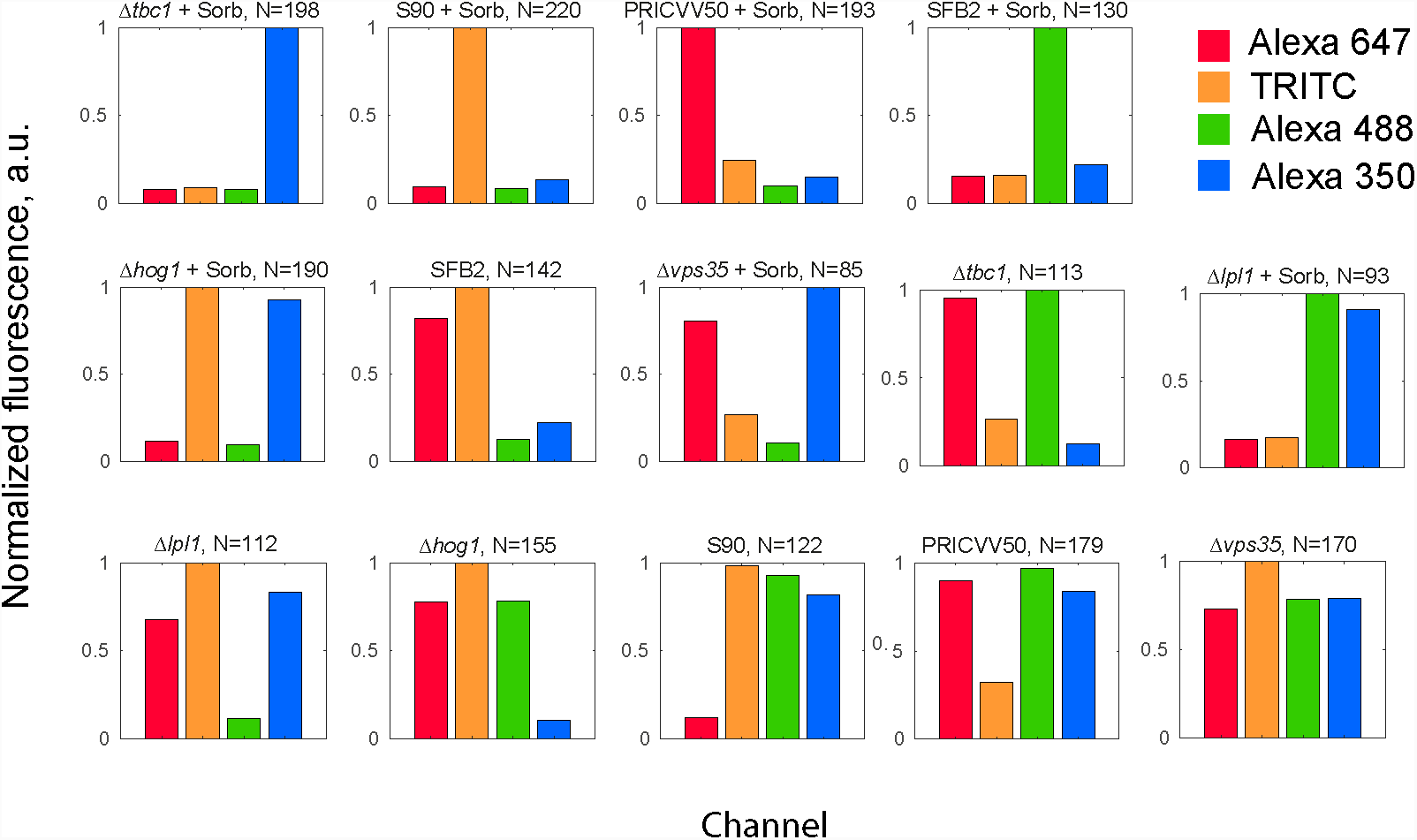
Barcode determination. The result of k-means classification of normalized fluorescence intensities measured in the osmotic shock experiment, showing all 14 combinations of fluorophores used in the experiment assigned to corresponding experimental condition.

**Fig. S4.**
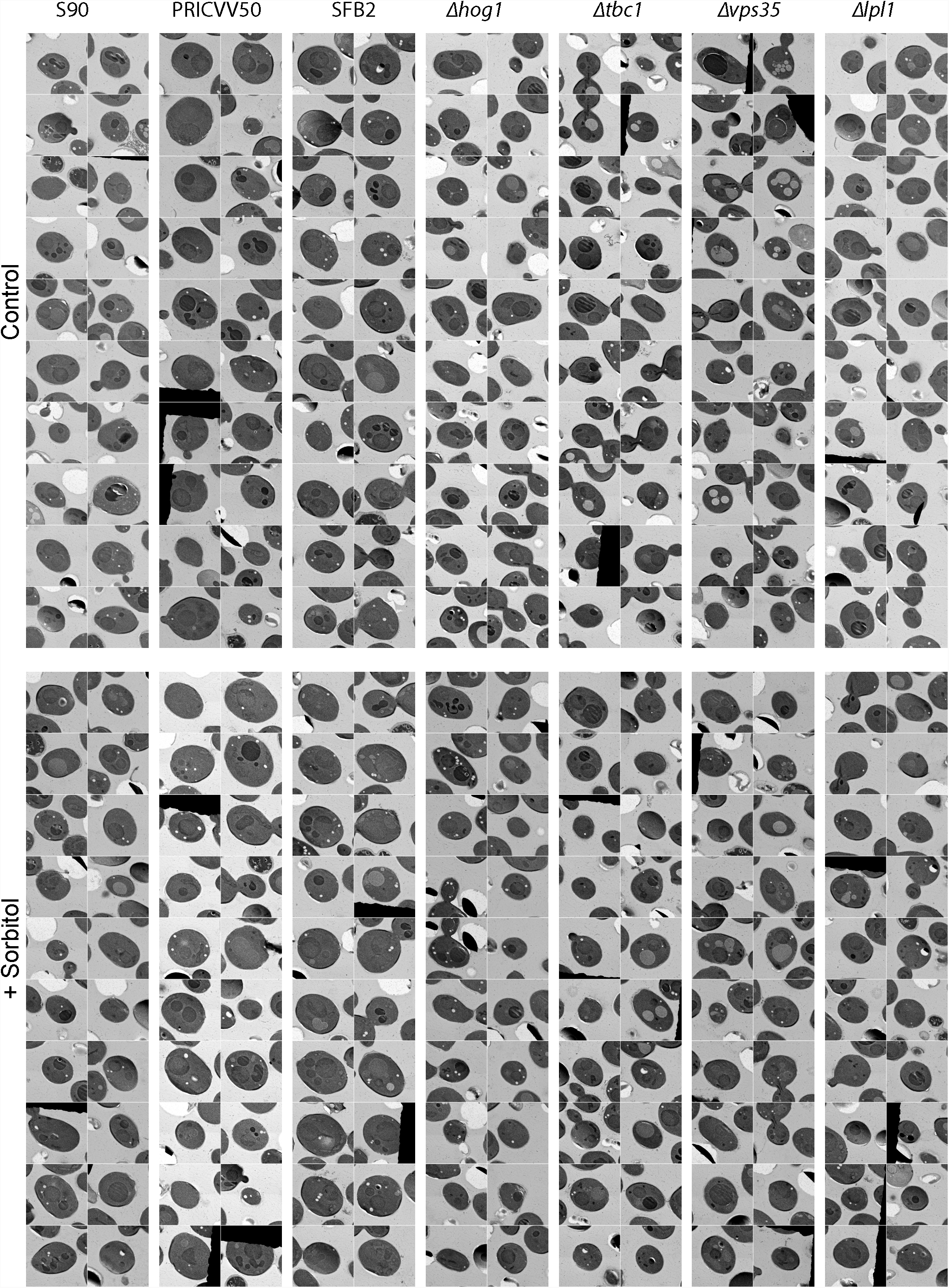
Gallery of yeast cell sections from different strains and experimental conditions. Twenty random examples of images from the final high-resolution EM dataset are shown for each strain. All cells display good preservation. A larger maximal cell cross-section area is apparent for wine yeast. Size of one panel is 8 µm.

**Fig. S5.**
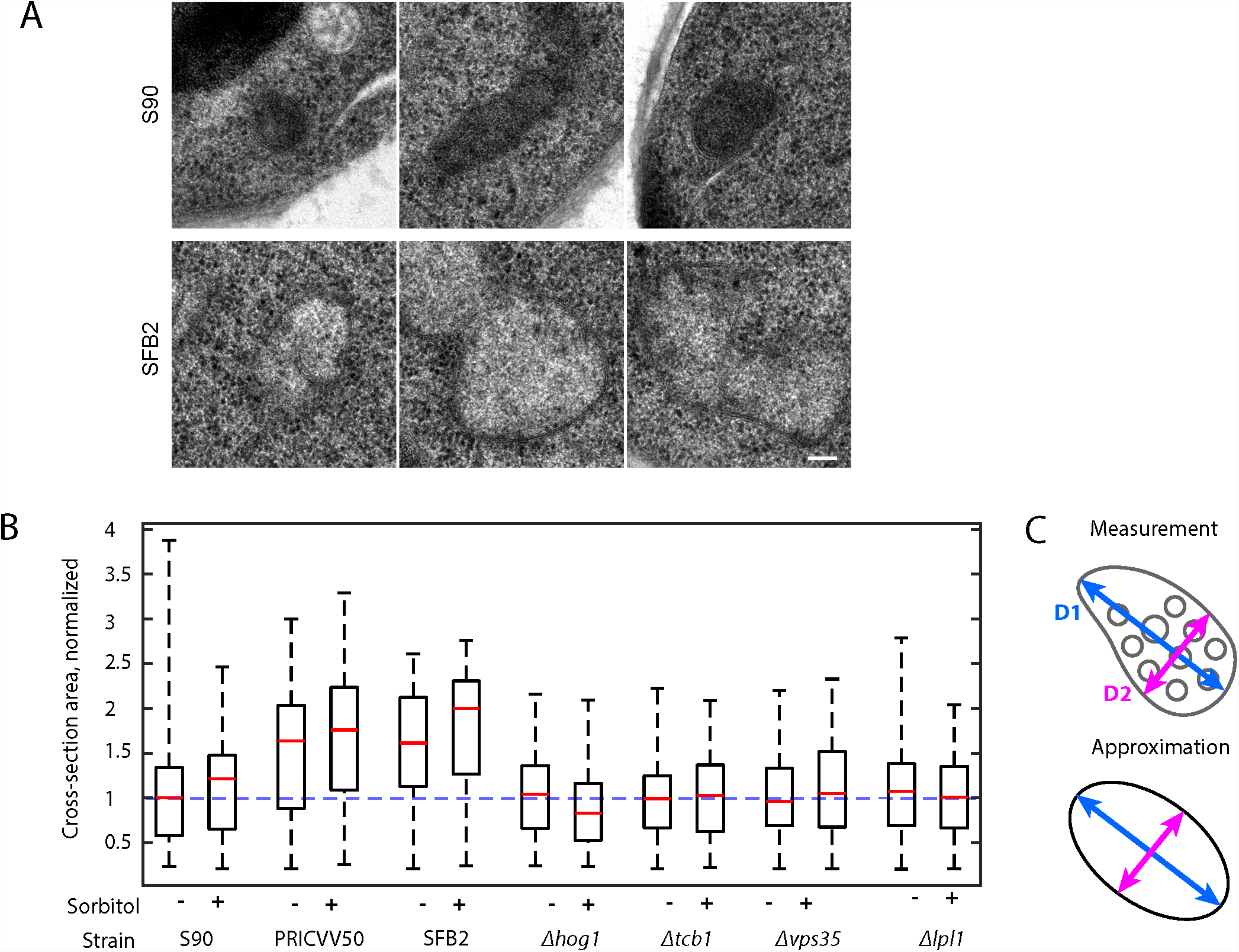
Mitochondria ultrastructure variation and MVB volume fraction measurement in wine and laboratory yeast strains. (A) Examples of mitochondria with normal morphology from the S90 strain (top) and abnormal swollen mitochondria from the SFB2 wine yeast (bottom). (B) Distributions of total mean cell cross-section areas for different strains and conditions, normalized to the S90 control (blue dashed line); boxes show the 25th, 50th, and 75th percentiles, whiskers show the range. PRICVV50 and SFB2 strains were characterized by increased cross-section areas, which signifies larger cells. PRICVV50 is a diploid (Novo et al., 2009), which explains their larger cell size, suggesting that SFB2, which has a similar size distribution is also a diploid. (C) The area of MVB cross-sections was measured not by point counting as in regular stereology but by approximation with an ellipse of similar shape: MVB major and minor axes (D1, D2) were determined (top) and area and circumference of an ellipse with the same axes were calculated (bottom) and used as an approximation of cross-section area and circumference of the measured MVB. See Materials and Methods for details. Scale bar is 100 nm.

**Fig. S6.**
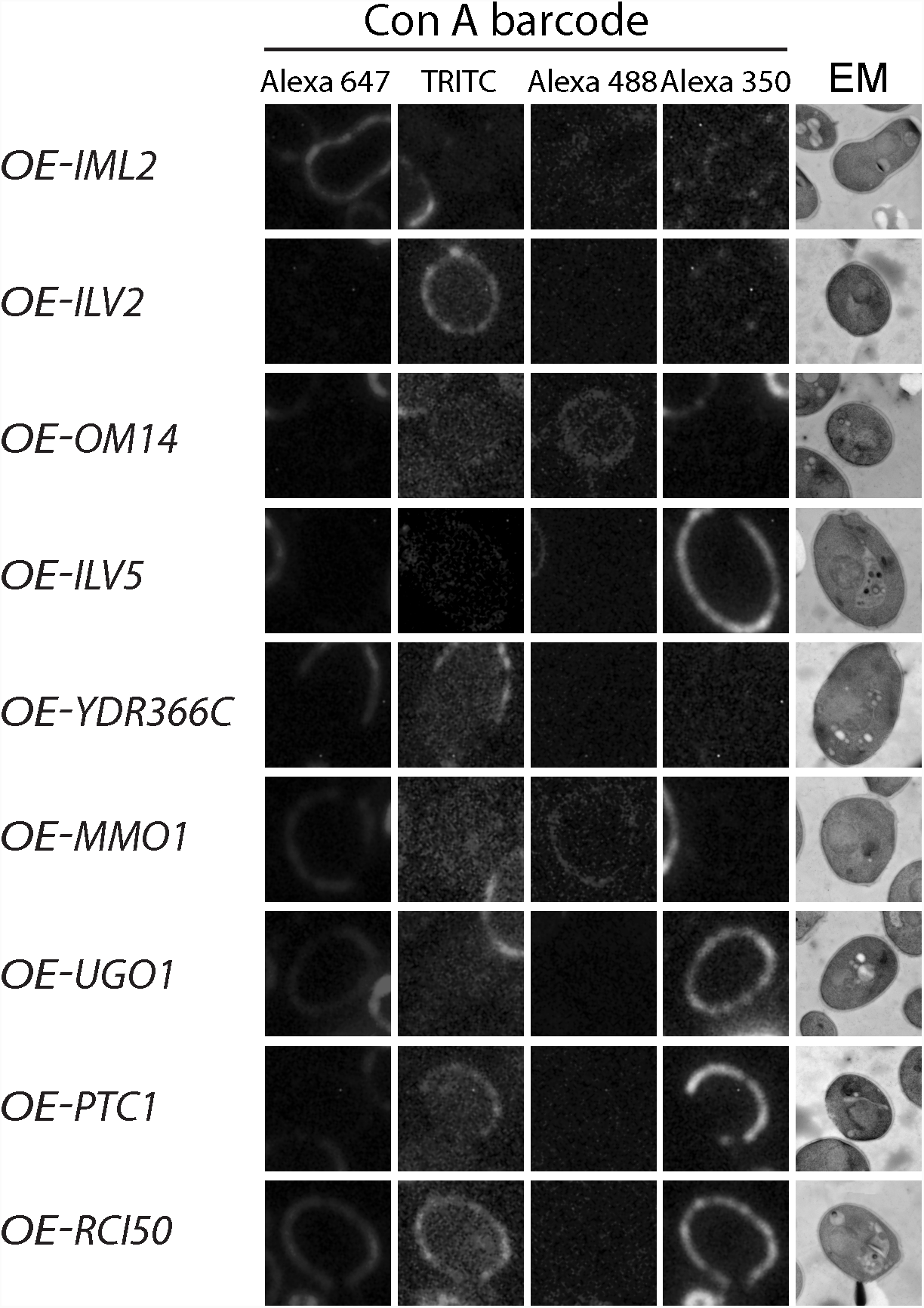
Examples of Con A barcodes for the mitochondria morphology experiment. First four columns show the LM images of the cell in different channels, fifth column shows EM image of the same cell. The name of the overexpressed (OE) protein is shown on the left for each strain. All strains are Δdnm1.

**Fig. S7.**
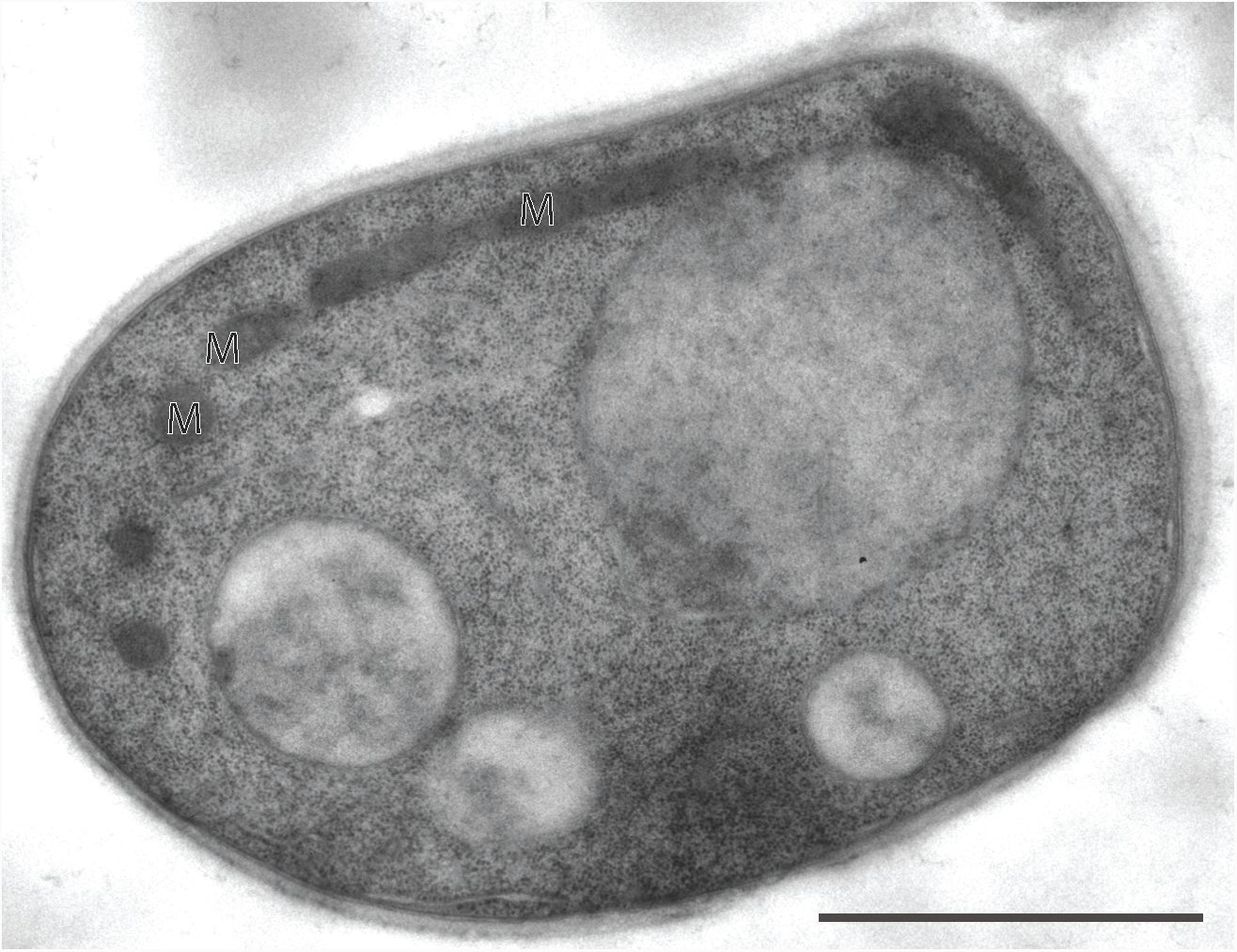
Morphology of the WT strain used in the mitochondrial ultrastructure study. Mitochondria are marked with “M”. Scale bar 1 µm.

**Fig. S8.**
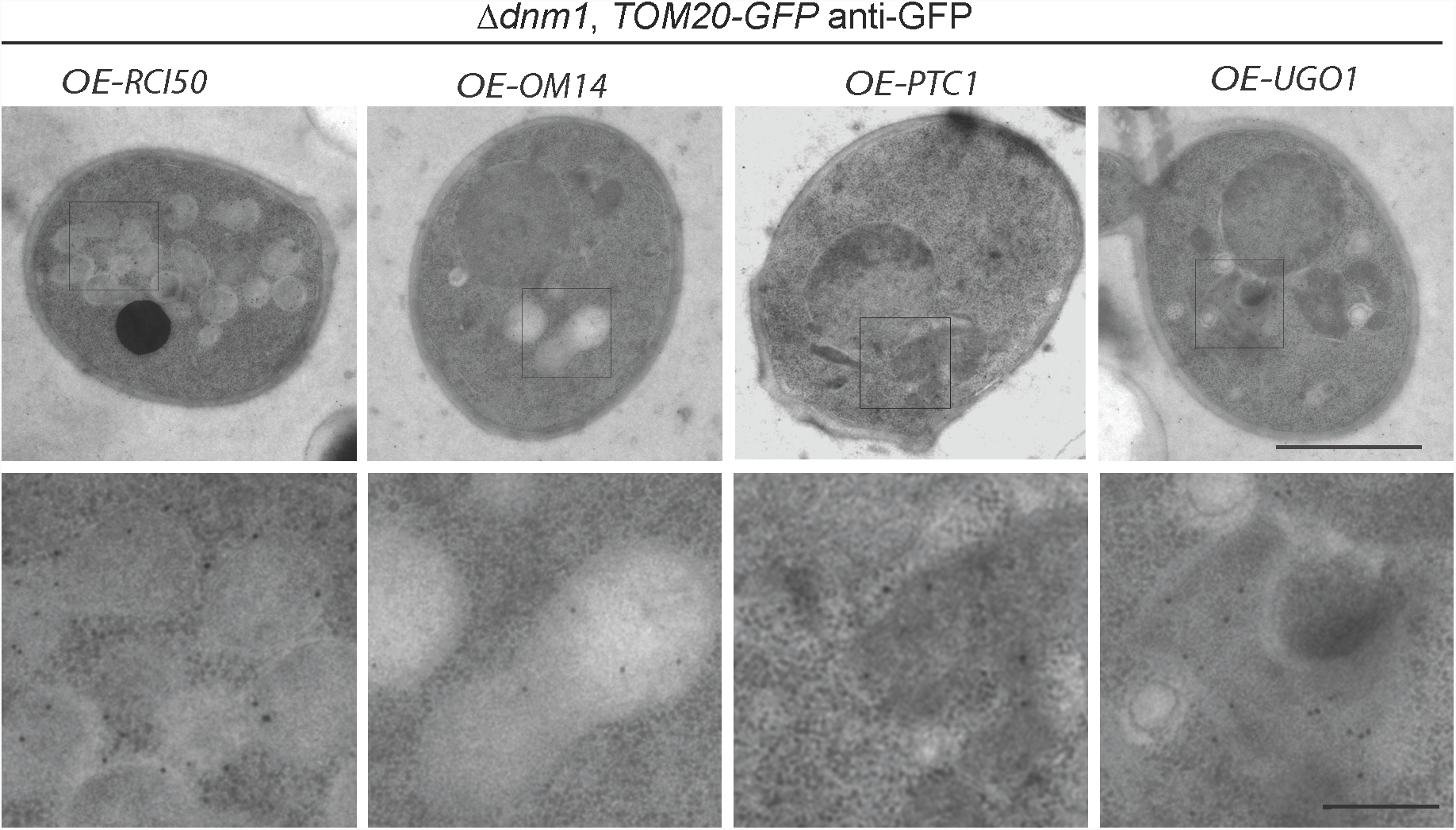
Using immunogold labeling for identification of the mitochondria in the MultiCLEM experiment. Three examples of Δdnm1 strains expressing TOM20-GFP and overexpressing proteins Ykl133c, Om14, and Ugo1 are selected from the grid labeled with anti-GFP antibodies. Top row – full cell view, bottom row – close up view of the regions marked by rectangles, showing immunogold labelling. Based on texture, bounding membrane morphology and labeling pattern, compartments labeled in the *OE-OM14* and *OE-UGO1* strains are likely to be vacuoles, while the structures in *OE-RCI50* may be mitochondria with an unusual morphology. Scale bars 1 µm (top) and 200 nm (bottom).

**Table S1.**
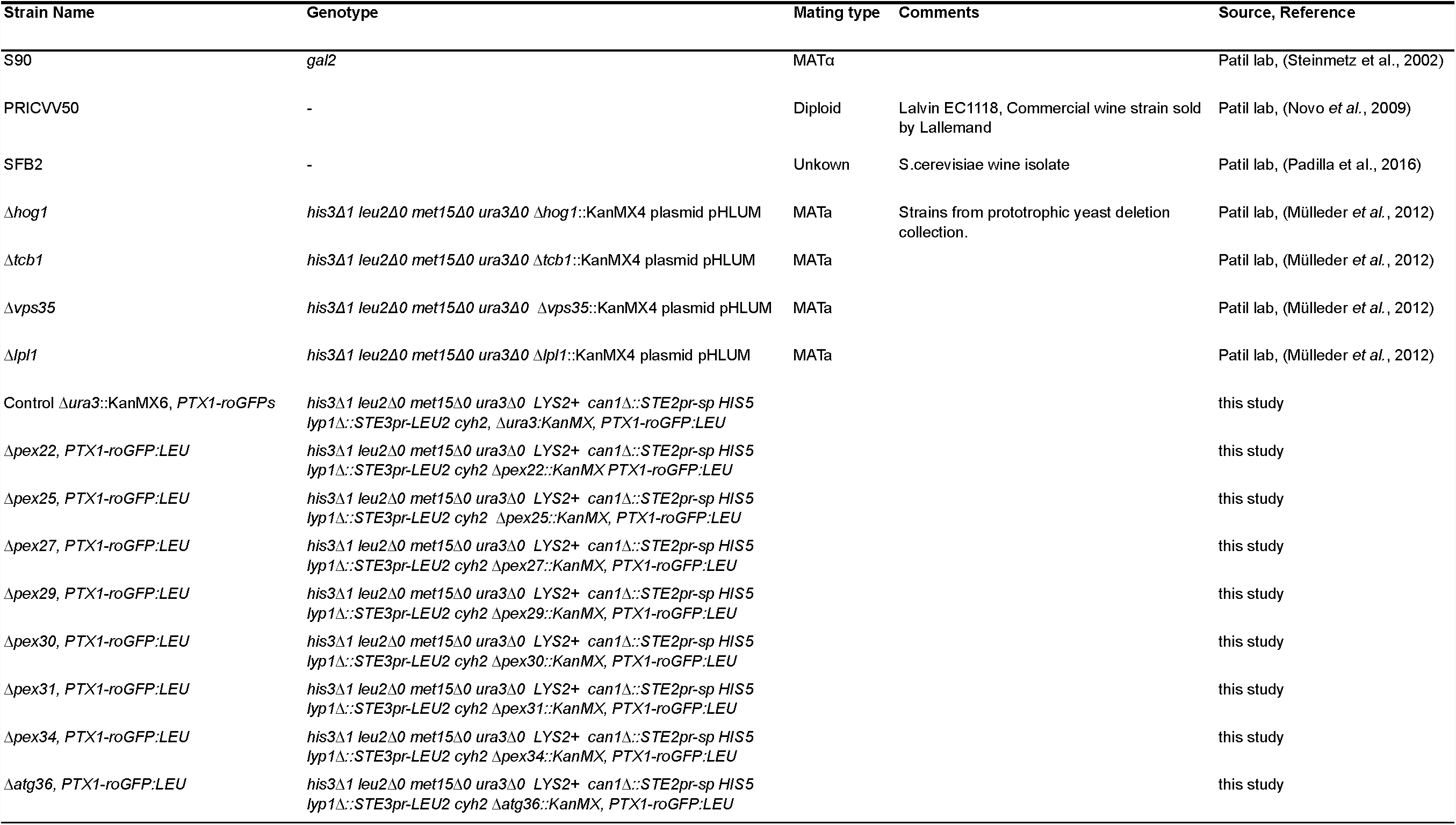
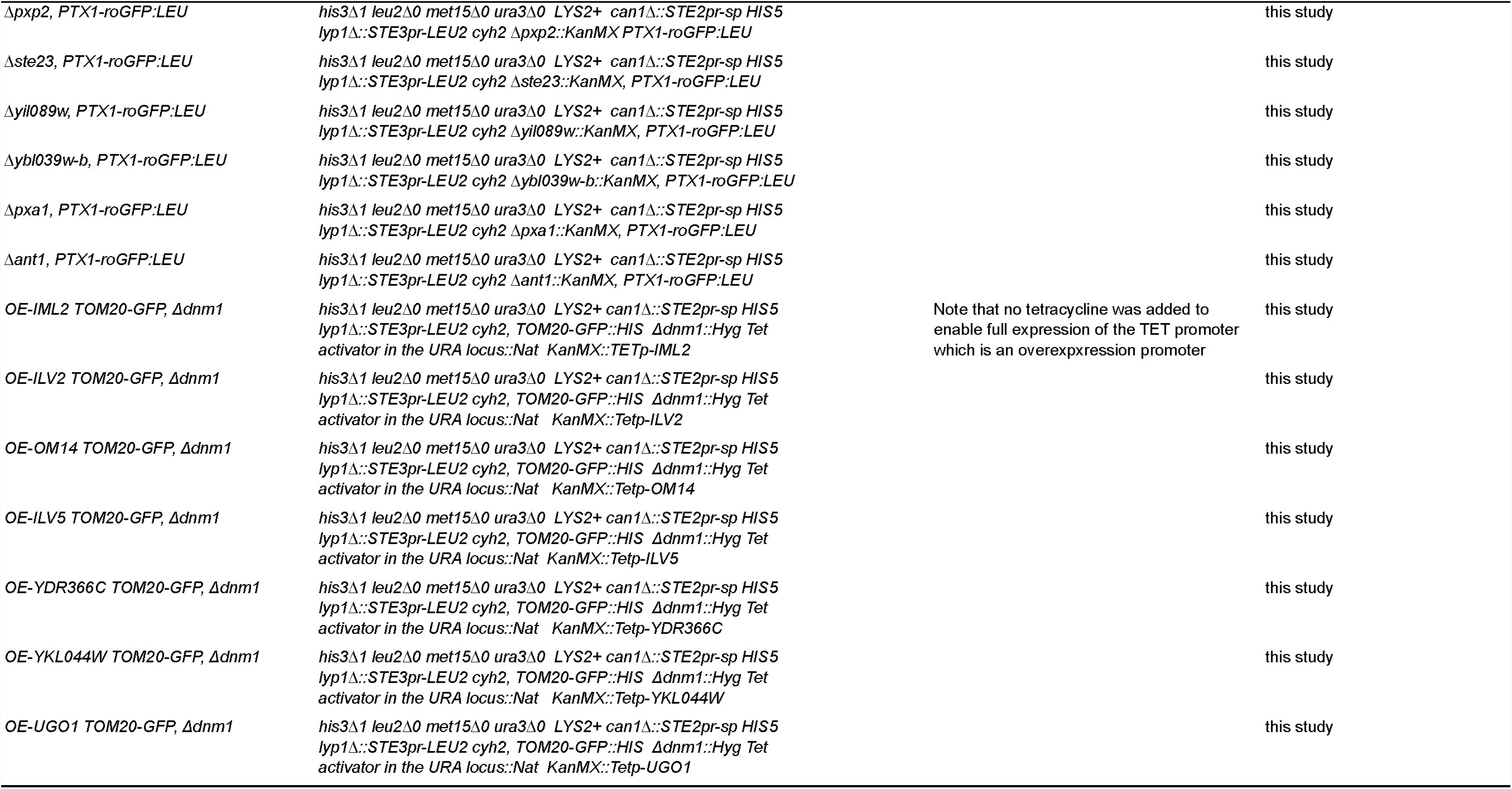
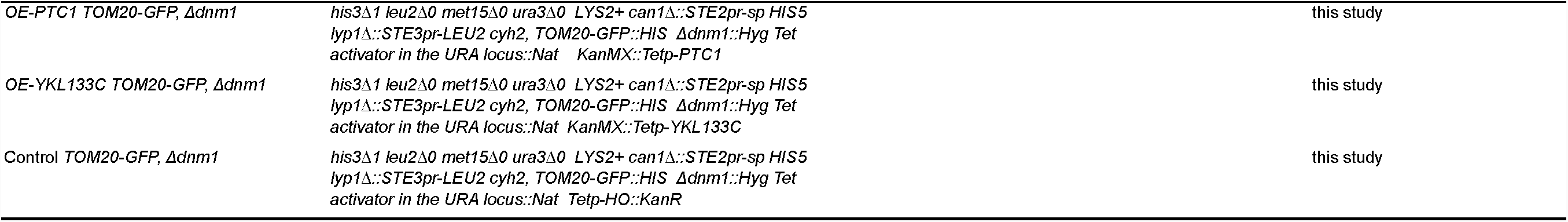
A list of all yeast strains used in this study as an excel (.xls) file.

| Strain Name | Genotype | Mating type | Comments | Source, Reference |
| --- | --- | --- | --- | --- |
| S90 | *gal2* | MATα | | Patil lab, (Steinmetz et al., 2002) |
| PRICVV50 | - | Diploid | Lalvin EC1118, Commercial wine strain sold by Lallemmand | Patil lab, (Novo *et al.*, 2009) |
| SFB2 | - | Unkown | *S.cerevisiae* wine isolate | Patil lab, (Padilla et al., 2016) |
| $\Delta hog1$ | *his3Δ1 leu2Δ0 met15Δ0 ura3Δ0 $\Delta hog1::$KanMX4 plasmid pHLUM* | MATa | Strains from prototrophic yeast deletion collection. | Patil lab, (Mülleder *et al.*, 2012) |
| $\Delta tcb1$ | *his3Δ1 leu2Δ0 met15Δ0 ura3Δ0 $\Delta tcb1::$KanMX4 plasmid pHLUM* | MATa | | Patil lab, (Mülleder *et al.*, 2012) |
| $\Delta vps35$ | *his3Δ1 leu2Δ0 met15Δ0 ura3Δ0 $\Delta vps35::$KanMX4 plasmid pHLUM* | MATa | | Patil lab, (Mülleder *et al.*, 2012) |
| $\Delta lpl1$ | *his3Δ1 leu2Δ0 met15Δ0 ura3Δ0 $\Delta lpl1::$KanMX4 plasmid pHLUM* | MATa | | Patil lab, (Mülleder *et al.*, 2012) |
| Control $\Delta ura3::$KanMX6, PTX1-roGFPs | *his3Δ1 leu2Δ0 met15Δ0 ura3Δ0 LYS2+ can1$\Delta::$STE2pr-sp HIS5 lyp1$\Delta::$STE3pr-LEU2 cyh2, $\Delta ura3::$KanMX, PTX1-roGFP:LEU* | | | this study |
| $\Delta pex22$, PTX1-roGFP:LEU | *his3Δ1 leu2Δ0 met15Δ0 ura3Δ0 LYS2+ can1$\Delta::$STE2pr-sp HIS5 lyp1$\Delta::$STE3pr-LEU2 cyh2 $\Delta pex22::$KanMX PTX1-roGFP:LEU* | | | this study |
| $\Delta pex25$, PTX1-roGFP:LEU | *his3Δ1 leu2Δ0 met15Δ0 ura3Δ0 LYS2+ can1$\Delta::$STE2pr-sp HIS5 lyp1$\Delta::$STE3pr-LEU2 cyh2 $\Delta pex25::$KanMX, PTX1-roGFP:LEU* | | | this study |
| $\Delta pex27$, PTX1-roGFP:LEU | *his3Δ1 leu2Δ0 met15Δ0 ura3Δ0 LYS2+ can1$\Delta::$STE2pr-sp HIS5 lyp1$\Delta::$STE3pr-LEU2 cyh2 $\Delta pex27::$KanMX, PTX1-roGFP:LEU* | | | this study |
| $\Delta pex29$, PTX1-roGFP:LEU | *his3Δ1 leu2Δ0 met15Δ0 ura3Δ0 LYS2+ can1$\Delta::$STE2pr-sp HIS5 lyp1$\Delta::$STE3pr-LEU2 cyh2 $\Delta pex29::$KanMX, PTX1-roGFP:LEU* | | | this study |
| $\Delta pex30$, PTX1-roGFP:LEU | *his3Δ1 leu2Δ0 met15Δ0 ura3Δ0 LYS2+ can1$\Delta::$STE2pr-sp HIS5 lyp1$\Delta::$STE3pr-LEU2 cyh2 $\Delta pex30::$KanMX, PTX1-roGFP:LEU* | | | this study |
| $\Delta pex31$, PTX1-roGFP:LEU | *his3Δ1 leu2Δ0 met15Δ0 ura3Δ0 LYS2+ can1$\Delta::$STE2pr-sp HIS5 lyp1$\Delta::$STE3pr-LEU2 cyh2 $\Delta pex31::$KanMX, PTX1-roGFP:LEU* | | | this study |
| $\Delta pex34$, PTX1-roGFP:LEU | *his3Δ1 leu2Δ0 met15Δ0 ura3Δ0 LYS2+ can1$\Delta::$STE2pr-sp HIS5 lyp1$\Delta::$STE3pr-LEU2 cyh2 $\Delta pex34::$KanMX, PTX1-roGFP:LEU* | | | this study |
| $\Delta atg36$, PTX1-roGFP:LEU | *his3Δ1 leu2Δ0 met15Δ0 ura3Δ0 LYS2+ can1$\Delta::$STE2pr-sp HIS5 lyp1$\Delta::$STE3pr-LEU2 cyh2 $\Delta atg36::$KanMX, PTX1-roGFP:LEU* | | | this study |

Table S1 continues on the next page
|  |  |  |  |
| --- | --- | --- | --- |
| *Δpxp2*, PTX1-roGFP:LEU | *his3Δ1 leu2Δ0 met15Δ0 ura3Δ0 LYS2+ can1Δ::STE2pr-sp HIS5 lyp1Δ::STE3pr-LEU2 cyh2 Δpxp2::KanMX PTX1-roGFP:LEU* |  | this study |
| *Δste23*, PTX1-roGFP:LEU | *his3Δ1 leu2Δ0 met15Δ0 ura3Δ0 LYS2+ can1Δ::STE2pr-sp HIS5 lyp1Δ::STE3pr-LEU2 cyh2 Δste23::KanMX, PTX1-roGFP:LEU* |  | this study |
| *Δyil089w*, PTX1-roGFP:LEU | *his3Δ1 leu2Δ0 met15Δ0 ura3Δ0 LYS2+ can1Δ::STE2pr-sp HIS5 lyp1Δ::STE3pr-LEU2 cyh2 Δyil089w::KanMX, PTX1-roGFP:LEU* |  | this study |
| *Δybl039w-b*, PTX1-roGFP:LEU | *his3Δ1 leu2Δ0 met15Δ0 ura3Δ0 LYS2+ can1Δ::STE2pr-sp HIS5 lyp1Δ::STE3pr-LEU2 cyh2 Δybl039w-b::KanMX, PTX1-roGFP:LEU* |  | this study |
| *Δpxa1*, PTX1-roGFP:LEU | *his3Δ1 leu2Δ0 met15Δ0 ura3Δ0 LYS2+ can1Δ::STE2pr-sp HIS5 lyp1Δ::STE3pr-LEU2 cyh2 Δpxa1::KanMX, PTX1-roGFP:LEU* |  | this study |
| *Δant1*, PTX1-roGFP:LEU | *his3Δ1 leu2Δ0 met15Δ0 ura3Δ0 LYS2+ can1Δ::STE2pr-sp HIS5 lyp1Δ::STE3pr-LEU2 cyh2 Δant1::KanMX, PTX1-roGFP:LEU* |  | this study |
| OE-IML2 TOM20-GFP, *Δdnm1* | *his3Δ1 leu2Δ0 met15Δ0 ura3Δ0 LYS2+ can1Δ::STE2pr-sp HIS5 lyp1Δ::STE3pr-LEU2 cyh2, TOM20-GFP::HIS Δdnm1::Hyg Tet activator in the URA locus::Nat KanMX::TETp-IML2* | Note that no tetracycline was added to enable full expression of the TET promoter which is an overexpression promoter | this study |
| OE-ILV2 TOM20-GFP, *Δdnm1* | *his3Δ1 leu2Δ0 met15Δ0 ura3Δ0 LYS2+ can1Δ::STE2pr-sp HIS5 lyp1Δ::STE3pr-LEU2 cyh2, TOM20-GFP::HIS Δdnm1::Hyg Tet activator in the URA locus::Nat KanMX::Tetp-ILV2* |  | this study |
| OE-OM14 TOM20-GFP, *Δdnm1* | *his3Δ1 leu2Δ0 met15Δ0 ura3Δ0 LYS2+ can1Δ::STE2pr-sp HIS5 lyp1Δ::STE3pr-LEU2 cyh2, TOM20-GFP::HIS Δdnm1::Hyg Tet activator in the URA locus::Nat KanMX::Tetp-OM14* |  | this study |
| OE-ILV5 TOM20-GFP, *Δdnm1* | *his3Δ1 leu2Δ0 met15Δ0 ura3Δ0 LYS2+ can1Δ::STE2pr-sp HIS5 lyp1Δ::STE3pr-LEU2 cyh2, TOM20-GFP::HIS Δdnm1::Hyg Tet activator in the URA locus::Nat KanMX::Tetp-ILV5* |  | this study |
| OE-YDR366C TOM20-GFP, *Δdnm1* | *his3Δ1 leu2Δ0 met15Δ0 ura3Δ0 LYS2+ can1Δ::STE2pr-sp HIS5 lyp1Δ::STE3pr-LEU2 cyh2, TOM20-GFP::HIS Δdnm1::Hyg Tet activator in the URA locus::Nat KanMX::Tetp-YDR366C* |  | this study |
| OE-YKL044W TOM20-GFP, *Δdnm1* | *his3Δ1 leu2Δ0 met15Δ0 ura3Δ0 LYS2+ can1Δ::STE2pr-sp HIS5 lyp1Δ::STE3pr-LEU2 cyh2, TOM20-GFP::HIS Δdnm1::Hyg Tet activator in the URA locus::Nat KanMX::Tetp-YKL044W* |  | this study |
| OE-UGO1 TOM20-GFP, *Δdnm1* | *his3Δ1 leu2Δ0 met15Δ0 ura3Δ0 LYS2+ can1Δ::STE2pr-sp HIS5 lyp1Δ::STE3pr-LEU2 cyh2, TOM20-GFP::HIS Δdnm1::Hyg Tet activator in the URA locus::Nat KanMX::Tetp-UGO1* |  | this study |

Table S1 continues
|  |  |  |
| --- | --- | --- |
| OE-PTC1 TOM20-GFP, Δdnm1 | *his3Δ1 leu2Δ0 met15Δ0 ura3Δ0 LYS2+ can1Δ::STE2pr-sp HIS5 lyp1Δ::STE3pr-LEU2 cyh2, TOM20-GFP::HIS Δdnm1::Hyg Tet activator in the URA locus::Nat KanMX::Tetp-PTC1* | this study |
| OE-YKL133C TOM20-GFP, Δdnm1 | *his3Δ1 leu2Δ0 met15Δ0 ura3Δ0 LYS2+ can1Δ::STE2pr-sp HIS5 lyp1Δ::STE3pr-LEU2 cyh2, TOM20-GFP::HIS Δdnm1::Hyg Tet activator in the URA locus::Nat KanMX::Tetp-YKL133C* | this study |
| Control TOM20-GFP, Δdnm1 | *his3Δ1 leu2Δ0 met15Δ0 ura3Δ0 LYS2+ can1Δ::STE2pr-sp HIS5 lyp1Δ::STE3pr-LEU2 cyh2, TOM20-GFP::HIS Δdnm1::Hyg Tet activator in the URA locus::Nat Tetp-HO::KanR* | this study |

**Table S2.**
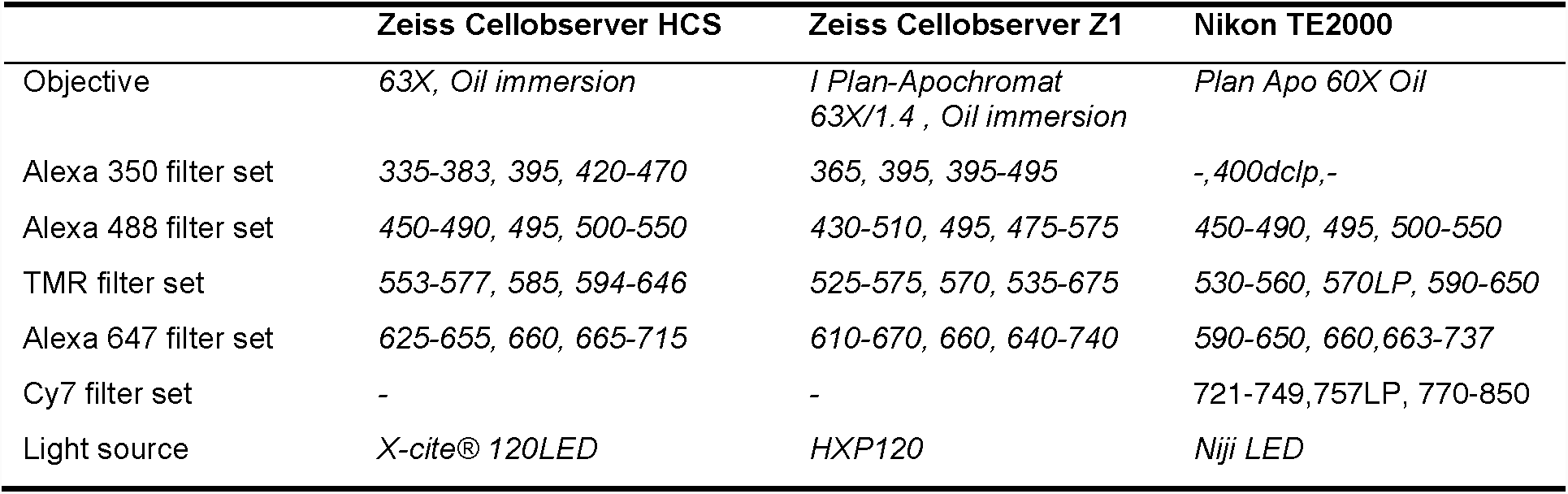
Fluorescent microscopes and filters used in this study. For each filter set, excitation filter transmission range, dichroic mirror splitting wavelength and emission filter transmission range are given in nm.

## Supplementary Data 1

Yury. S. Bykov, Nir Cohen et al.

### The yeast fluorescent barcoding protocol

#### Materials and equipment

- Yeast media
- Phosphate buffered saline (PBS)
- Concanavalin A (Con A) storage stock solutions, 2.5 mg/ml in PBS. Prepare from commercially available lyophilized powder, sonicate to dissolve, aliquot and store at −20°C only. 50-100 µl aliquots are usually a good choice.
- Tubes or plates with gas-permeable membrane
- Spectrophotometric cuvettes (plastic)
- Fresh YPAD agar plate
- Razor blade
- 0.3 mm and 0.1/0.2 mm aluminum membrane carriers for HPM010
- Tweezers and forceps
- Small flat metal spatula (0.5 cm wide)
- Pipette (0.5-10 μl)
- MILLIPORE filtering setup with 0.45 um filter
- Spectrophotometer
- Yeast incubator
- Ultrasonic bath
- Centrifuge for plates (e.g. Eppendorf 5820R)
- Standard benchtop centrifuge for 1.5 ml tubes
- (Optional) Ultracentrifuge and rotor, 1.5 ml ultracentrifuge tubes (e.g. Beckman Optima Max with TLA55 rotor)
- HPM010 high-pressure freezing machine

#### Yeast cultures

For freezing 3-5 HPM010 carriers the total amount of yeast cells should be 15-20 ml at OD_600_=0.6. This means that for a 15-fold multiplexed experiment each strain should be grown in 1 ml to OD_600_=0.6. This can be done in separate tubes or in a 24-well plate. Separate tubes are more difficult to handle but easier to grow (in a normal incubator). 24-well plates require smaller amplitude and higher speed orbital shaker (they cannot be properly mixed with a normal incubator for flasks). One can do it by using a plate shaker, or by sticking a plate to the top of the Eppendorf Thermomixer which is in turn placed inside a regular incubator at 30°C. The cultures can be prepared the day before to reach the OD=0.6 in the morning, so staining can be started immediately, or grown from diluted overnight cultures during the day. We use the following growth protocol to achieve desired cell density before freezing in the afternoon.

1. Dilute the overnight cultures in tubes or in a plate. To get OD=0.6 at 14:00 one would need an OD of approximately 0.15 at 09:30 for relatively fast growing strains.
2. If using a plate, seal it with a gas-permeable membrane, and install on the thermomixer or plate shaker inside the incubator at 30°C. The shaking speed should be 500-600 rpm.

### Staining solution preparation

Staining solutions are prepared in two steps to reduce the errors and speed up the process. First concentrated 5X Con A stocks are prepared from the frozen Con A storage stocks. Relative brightness of different fluorescent channels can be adjusted at this stage. Then 5X stocks are mixed in equal amounts to produce the final staining solutions.

1. Thaw Con A storage stock solutions (2.5 mg/ml in PBS) and sonicate for 5 minutes in an ultrasonic bath to dissolve the aggregates.
2. Spin down any remaining large aggregates on a tabletop centrifuge at maximum speed, (16000 rcf), 4°C.
3. Prepare 5X stock solutions for fast mixing. To compensate for different conjugate brightness the concentrations can be adjusted at this stage (Table 1).
4. Optional: ultracentrifuge 5X stocks 30 min at 200000 g (45000 rpm) on TLA55 rotor of Beckmann tabletop centrifuge to remove more aggregates.
5. Mix the 5X stocks with each other and with PBS according to the pattern shown in Table 2 to get final staining solutions^1^. To increase speed and accuracy of mixing, label the tube rims in advance using markers of different colors showing which 5X stocks are added to which tube. First add the blue 5X stock to all tubes with the blue mark, then the green 5X stock to all tubes with the green mark etc.

**Table 1.** Example of 5X Con A stocks preparation recipe. The amount of Con A storage stock can be easily increased if it starts going bad, or more labelling intensity is needed without adjusting the final staining solutions preparation.

| Conjugate name | Con A storage stock, µl | PBS, µl | Total, µl |
| --- | --- | --- | --- |
| **B: Con A Alexa 350** | 120 | 930 | 1050 |
| **G: Con A Alexa 488** | 40 | 1010 | 1050 |
| **O: Con A TMR 555** | 50 | 1000 | 1050 |
| **R: Con A Alexa 647** | 100 | 950 | 1050 |
| **IR: Con A Cy7 (home made)** | 250 | 800 | 1050 |

<sup>1</sup> The scheme can easily be extended to 5 colors with the same total volume – 600 µl – by adding extra 120 µl of the fifth stock solution and reducing the amount of PBS accordingly.

**Table 2.** Final staining solutions preparation. For each staining solution the amount of each 5X stock (denoted by the first letter of color, see Table 1) and PBS to add is indicated in µl.

| Solution | B | G | O | R | PBS | Solution | B | G | O | R | PBS |
| --- | --- | --- | --- | --- | --- | --- | --- | --- | --- | --- | --- |
| 1 | 120 | 0 | 0 | 0 | 480 | 9 | 0 | 120 | 0 | 120 | 360 |
| 2 | 0 | 120 | 0 | 0 | 480 | 10 | 0 | 0 | 120 | 120 | 360 |
| 3 | 0 | 0 | 120 | 0 | 480 | 11 | 120 | 120 | 120 | 0 | 240 |
| 4 | 0 | 0 | 0 | 120 | 480 | 12 | 120 | 120 | 0 | 120 | 240 |
| 5 | 120 | 120 | 0 | 0 | 360 | 13 | 120 | 0 | 120 | 120 | 240 |
| 6 | 120 | 0 | 120 | 0 | 360 | 14 | 0 | 120 | 120 | 120 | 240 |
| 7 | 120 | 0 | 0 | 120 | 360 | 15 | 120 | 120 | 120 | 120 | 120 |
| 8 | 0 | 120 | 120 | 0 | 360 |  |  |  |  |  |  |

### Staining and freezing

1. Check the OD_600_ of all or a representative subset of strains. It should be 0.5-0.6. When the right OD is achieved, start filling HPM010 with nitrogen before proceeding to the next steps. Filling takes 15-20 minutes.
2. Spin down the cells at 1500 rcf for 5 min.
3. Carefully remove the plate from the centrifuge and discard the supernatants without disturbing the pellets. The pellets are very loose at this stage! Immediately after removing the supernatant fill the well with the staining solution and mix well.
4. Make sure the pellets are resuspended well and put the plate on the mixer in the incubator for 5-10 minutes. Avoid exposing to bright light.
5. Remove the plate from the incubator and spin it down at 1500 rcf for 5 min. Discard the supernatants and resuspend pelleted cells in medium. The pellets will cover the whole well bottom and may require thorough mixing in order to collect and resuspend all of the cells.
6. During the resuspension or afterwards pool all resuspended pellets in one 50 ml falcon tube. Vortex it well.
7. Proceed immediately. Collect the yeast on a filter using the MILLIPORE filtering device, and immediately put the filter on a fresh YPAD plate. Make sure it is wetted by the plate and that there are no bubbles in-between the filter and the agar surface.
8. Scrape the cell slurry on the filter into a dense clump with a spatula. The cells should be as concentrated as possible, but not too much: the concentration is too high and cells need diluting if after placed into HPM010 carrier, the slurry droplet surface quickly becomes opaque. To make the slurry more dilute, press filter with a spatula to squeeze water from the plate. To concentrate the cells lift the filter for a short time to make the cells dry a little bit.
9. Cut off the end of the pipet tip to make the hole wider, pipette 1.5-1.75 μl of suspension into the tip, holding the cell clump with a spatula. Do not press too hard to avoid diluting the suspension.
10. Apply the suspension to the 0.1 mm cavity of a high-pressure freezer membrane carrier, cover with a flat side of a 0.3 mm cavity carrier, and freeze immediately.

## Footnotes

1 The scheme can easily be extended to 5 colors with the same total volume – 600 μl – by adding extra 120 μl of the fifth stock solution and reducing the amount of PBS accordingly.

